# Cross-species phenotypic profiling uncovers functional determinants of bacterial cold shock adaptation

**DOI:** 10.1101/2025.09.22.676666

**Authors:** Hao Fan, Andrew Briercheck, Vivek K Mutalik, Mengwen Xiong, Yan Zhang

## Abstract

Temperature shifts impose broad physiological stress, requiring precise and dynamic regulatory programs to restore cellular homeostasis. While the heat shock response is well characterized, the mechanisms underlying cold shock response (CSR) remain less understood. To identify genes critical for cold adaptation, we applied transposon sequencing (Tn-seq) to monitor mutant fitness across the full course of CSR and sustained low-temperature growth in two mesophilic bacteria, *Escherichia coli* and *Bacillus subtilis*. In *B. subtilis*, phenotypic profiling revealed a temporally structured program: membrane fluidity and cell wall remodeling were most critical in the early stage of CSR, whereas post-transcriptional regulation became essential during late-stage recovery to reprogram gene expression and restore growth. Cross-species comparison uncovered both conserved and species-specific mechanisms, with RNA metabolism and ribosome/translation regulators playing broad roles. Specifically, we identified a conserved synergy between two ribosomal RNA methyltransferases, RsmA and RsmH, in promoting cold adaptation. In *B. subtilis*, mutants lacking these enzymes exhibited significant delay in translation recovery following cold-induced global inhibition. Together, these findings provide a comparative, systems-level view of bacterial cold adaptation and establish a framework for exploring stress responses in pathogens and extremophiles.

## Introduction

Bacteria must adapt rapidly to environmental stresses, with temperature fluctuations presenting a constant challenge. For host-associated species, sudden transitions into or out of warm-blooded hosts can trigger regulatory programs influencing virulence and survival (1), while psychrotrophic bacteria adapted to near-freezing temperatures pose increasing risks to food safety and public health (2, 3). Unlike the well-characterized heat shock response, which is driven primarily by transcriptional regulation, the cold shock response (CSR) lacks a dedicated transcription factor. Instead, the CSR relies mainly on post-transcriptional processes, such as RNA stabilization, ribosome assembly, and translational control (4–6). However, the molecular mechanism underlying these regulators, and how post-transcriptional control broadly facilitates cold adaptation, remain poorly understood. This gap makes CSR a valuable model for uncovering fundamental principles of bacterial stress adaptation.

Most mechanistic insights into the CSR have come from studies in *Escherichia coli*. Following a downshift from 37°C to 10°C, *E. coli* undergoes a ∼6-hour lag phase before resuming slower growth at 10°C (7, 8). Early 2D-gel proteomics identified a small set of proteins strongly induced after cold shock despite the global suppression of protein synthesis (7). Later studies provided deeper characterization of these cold-inducible proteins, which include RNA chaperones such as cold shock proteins (Csps) (5, 9), RNA helicases (e.g., DeaD) (10), ribonucleases (e.g., RNase R, PNPase) (11, 12), translational factors (e.g., IF1, IF3, RbfA) (13–15), DNA topology modulators (e.g., DNA gyrase, H-NS) (16, 17), and membrane-modifying enzymes (e.g., LpxP) (18). A more recent multi-omics study revealed an mRNA surveillance system that drives global translation recovery during cold adaptation (6). Two groups of factors are central to this system: Csps, which act as RNA chaperons to resolve cold-disrupted mRNA secondary structures and broadly restore translation; and RNase R, which degrades highly folded mRNAs to facilitate post-transcriptional reprogramming. A negative feedback loop then fine-tunes Csp expression, enabling timely termination of the CSR once adaptation is achieved (6).

Despite extensive knowledge of individual cold-inducible factors in *E. coli*, the overall architecture of the CSR remains poorly defined, particularly how it progresses over time, how it varies between species, and which genes are truly critical for adaptation and survival. Cold shock perturbs numerous thermodynamically sensitive processes, yet the mechanism by which cells prioritize and coordinate these responses across CSR stages remains unclear. Cross-species comparisons are limited, and the functional contributions of most genes have not been systematically tested, leaving fundamental questions about the organization and evolution of CSR unresolved.

Here, we applied cross-species transposon sequencing (Tn-seq) to map the genetic requirements for cell fitness across distinct CSR stages and during continuous cold growth. We compared *E. coli* and *Bacillus subtilis*, two phylogenetically and ecologically distinct mesophiles that thrive at moderate temperatures (20-45°C) but exhibit markedly different growth behavior following temperature downshift. Genome-wide fitness profiling revealed a temporally structured response in *B. subtilis*, with distinct gene sets required at early and late stages of CSR as well as during long-term cold growth. *E. coli* undergoes a brief nongrowing acclimation phase, limiting the resolution of stage-specific requirements. Comparative analysis uncovered key cold adaptation factors that are either species-specific or broadly conserved, including two universal rRNA methyltransferases and multiple previously uncharacterized genes. Together, this work provides a comparative, time-solved, and systems-level view of cold adaptation in bacteria.

## Results

### Characterization of bacterial growth phenotypes after cold shock

Systematic characterization of cold shock growth phenotypes is limited outside *E. coli*. To address this, we compared the behavior of *E. coli* MG1655 and *B. subtilis* 168 strains after shifting exponentially growing cultures at 37°C to 18°C, 15°C, 12°C, or 9°C, and then maintaining them at the new temperature. These target temperatures span a range below the optimal growth window for both species (20–45°C) but remain above the lower growth limit for *E. coli* (∼7.5°C) (19).

In line with prior studies (7, 20), *E. coli* displayed a lag period (also known as the acclimation phase) immediately after cold shock, with its duration increasing as the temperature drop became more severe (Fig. 1A, B). The adapted growth rate also decreased with lower temperatures (Fig. 1C). In contrast, *B. subtilis* 168 lacked a defined no-growth acclimation phase: following a cold shock from 37°C to 18°C, cell growth gradually slowed over an extended period (∼16 hours) before becoming fully adapted (Fig. 1D), reaching a steady-state growth rate comparable to that of *E. coli* at 18°C (Fig. 1E). However, upon a temperature shift to 15°C, *B. subtilis* failed to fully resume growth over the following two days, with only subtle increase in cell density (Fig. 1D, S1A). Notably, cold shock to 12°C triggered cell lysis, which became more pronounced upon shifting to even lower temperatures (Fig. 1D).

**Figure 1.**
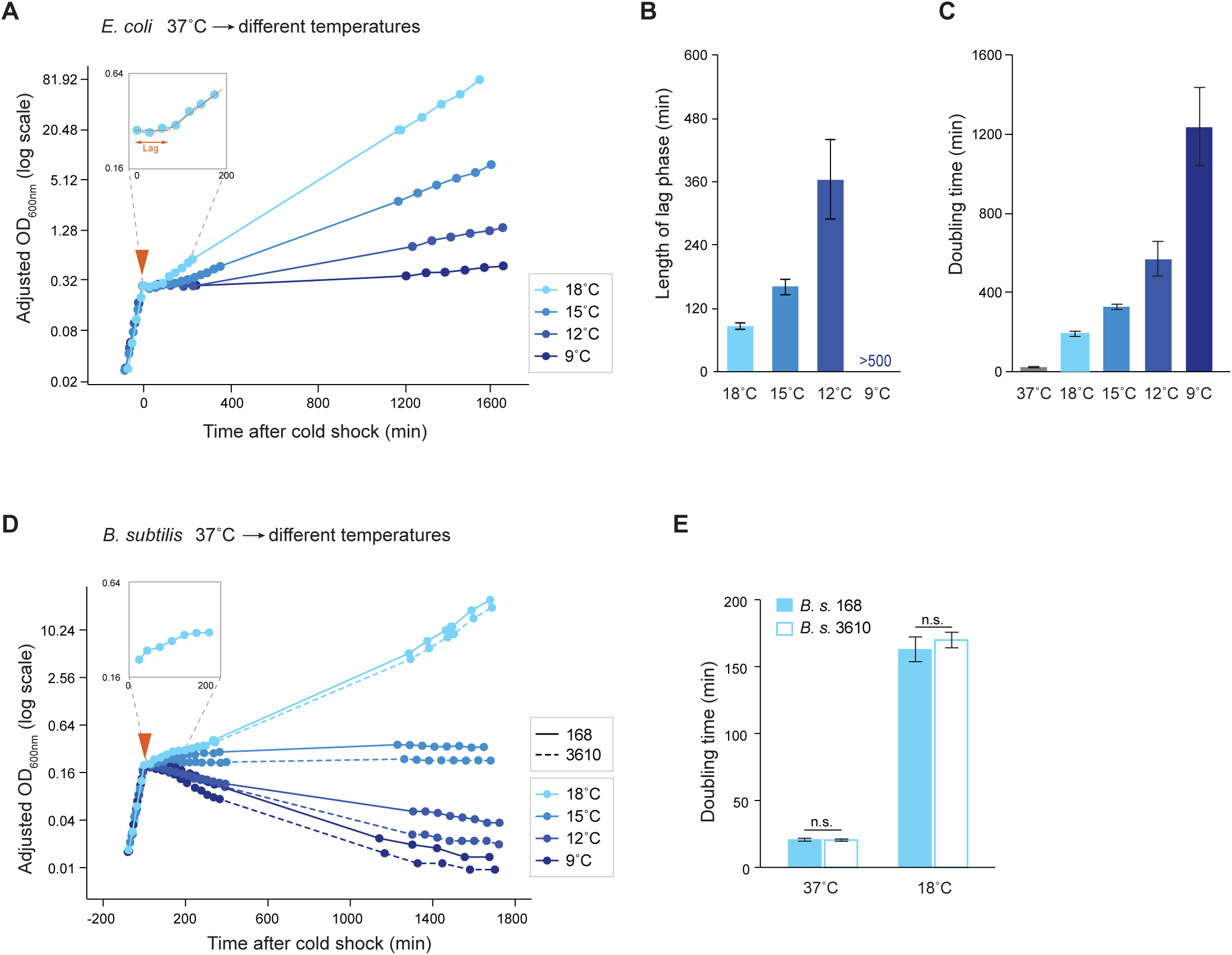
Growth phenotypes of *E. coli* and *B. subtilis* after cold shock to different temperatures. **(A)** Growth curve of *E. coli* MG1655 cells at 37°C and after shifting to 18°C, 15°C, 12°C, or 9°C. Cold shock was induced by mixing the 37°C culture with pre-chilled LB medium in defined volumes, allowing an immediate temperature shift and synchronized cold shock response across the population. X-axis: time after cold shock (min); Y-axis: adjusted optical density at 600nm (OD_600_) on a log scale. OD_600_ was adjusted at the transitions of cold shock and back-dilutions (see Methods) to allow continuous readings. The inset shows a zoomed-in view of the lag phase following cold shock before recovery to a new steady-state growth. All the growth curves shown in this and subsequent figures represent one of at least 3 biological replicates. **(B)** Bar graph showing the average duration of the lag phase (min) of *E. coli* MG1655 after cold shock from 37°C to various low temperatures. Error bars: standard deviation (n = 3). For the 9°C cold shock, the lag phase duration was too long to measure precisely, so an estimated value is shown. **(C)** Doubling times of *E. coli* MG1655 cells during steady-state growth at various temperatures, averaged from 3 biological replicates. Error bars: standard deviation. **(D)** Growth curves of *B. subtilis* 168 and 3610 strains at 37°C and after shifting to various low temperatures. X-axis: time after cold shock (min); Y-axis: adjusted OD_600_ on a log scale. OD_600_ was adjusted similarly to **(A).** The inset shows the continuous but slower growth (no immediate lag phase) following cold shock. **(E)** Average doubling time of *B. subtilis* 168 and 3610 strains during steady-state growth at various temperatures. Error bars: standard deviation (n = 3); n.s.: not significantly different based on student’s t-test.

To test whether these phenotypes are specific to the laboratory strain *B. subtilis* 168, we examined the undomesticated wild-type strain NCIB 3610 (21). Compared to 168, strain 3610 lysed even more rapidly when shifted to temperatures below 15°C, but showed similar steady-state growth at 18°C (Fig. 1D). These differences may reflect mutations acquired during laboratory domestication of strain 168. Together, these results indicate that *E. coli* and *B. subtilis* employ species-specific strategies to cope with cold shock, and that cellular adaptation is not determined solely by the magnitude of temperature downshift.

### Tn-seq enables cross-species comparison of key Cold Shock Response (CSR) factors

To compare cold adaptation strategies between *E. coli* and *B. subtilis*, we employed transposon sequencing (Tn-seq) to identify genes required for fitness during distinct CSR stages and continuous low-temperature growth using previously constructed libraries (22, 23) (Fig. 2). Gene-specific fitness values were calculated by comparing Tn-seq counts across each open reading frame (ORF) between adjacent time points, after normalizing for the number of generations (doublings) and correcting for chromosome position bias (Figs. 2A, S2-S4; see Methods) (24). Fitness values near 1 indicate little or no growth effect relative to wild type; values <1 indicate functional importance of the gene, and values >1 suggest a growth advantage upon gene disruption.

**Figure 2.**
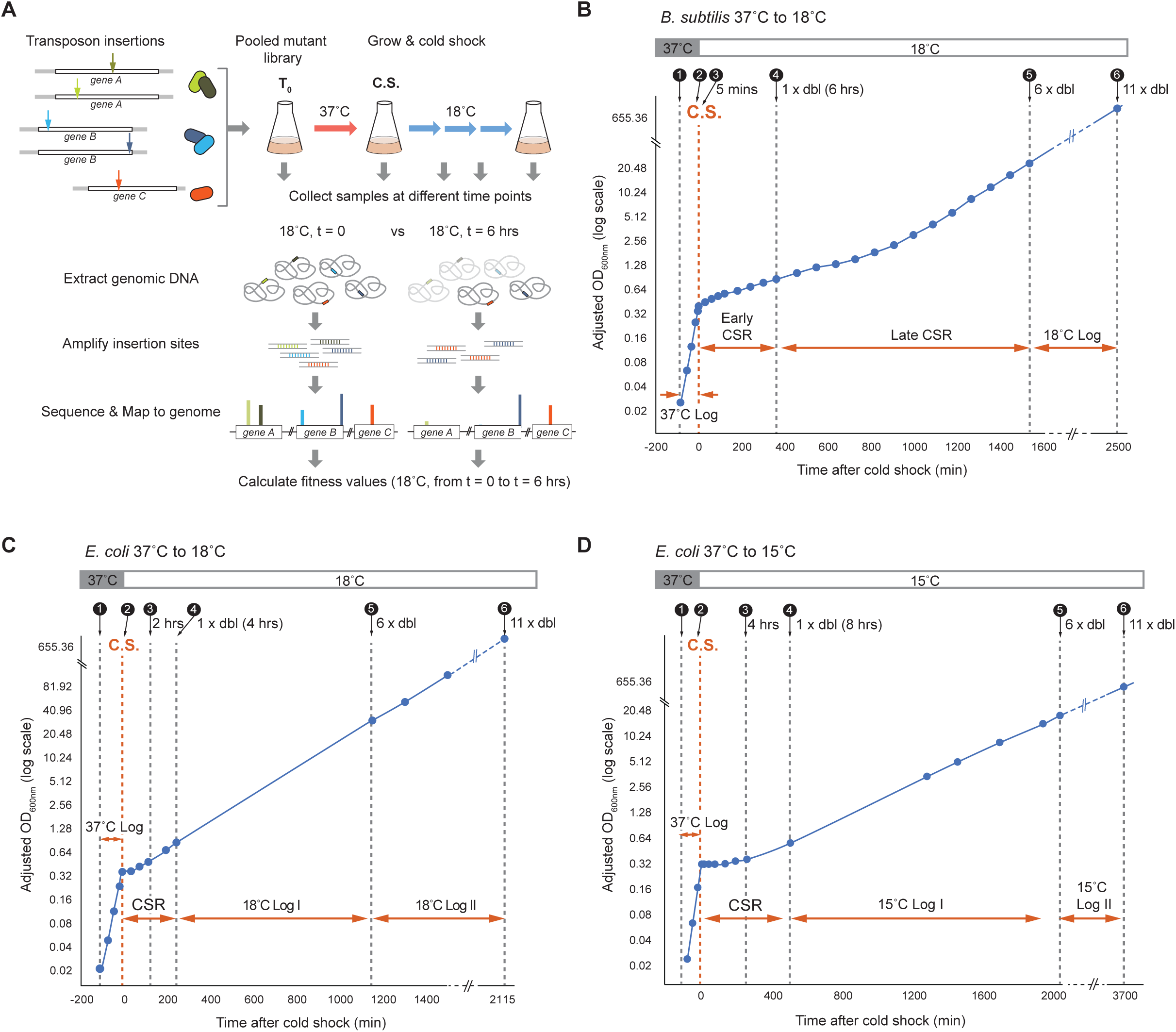
Transposon sequencing (Tn-seq) setup and data analysis. **(A)** Schematic overview of the Tn-seq experiments and fitness value calculation. Pooled transposon mutant libraries of *E. coli* or *B. subtilis* were grown at 37°C and then cold shocked (C.S.) to 18°C. Samples were collected at multiple time points before and after the temperature shift. Genomic DNA from each sample was used to amplify the transposon insertion sites (*B. subtilis*) or associated barcodes (*E. coli*), which were then sequenced to quantify the relative abundance of different transposon mutants. Fitness values for individual genes were calculated by comparing normalized Tn-seq signals between two time points (e.g., during the first 6 hours after cold shock, as shown). A decrease in Tn-seq read counts (fitness value < 1; e.g., *gene A*) indicates that the gene is functionally important during that time interval. **(B)** Timeline for *B. subtilis* Tn-seq sample collection. Samples 1–6: (1) t_0_ control, (2) 37°C before cold shock, (3) 5 minutes, (4) 1 doubling (1 x dbl), (5) 6 doublings (6 x dbl), and (6) 11 doublings (11 x dbl) after shifting to 18°C. Defined time intervals are indicated by red arrows, including 37°C log phase (37°C Log), early and late stages of cold shock response (Early CSR and Late CSR), and 18°C adapted phase (18°C Log). C.S.: cold shock. **(C, D)** Timeline for *E. coli* RB-Tnseq sample collection. Samples 1–6: (1) t_0_ control, (2) 37°C before cold shock, (3) 2 hours (18°C) or 4 hours (15°C), (4) 1 doubling (1 x dbl), (5) 6 doublings (6 x dbl), and (6) 11 doublings (11 x dbl) after shifting to 18°C **(C)** or 15°C **(D)**. Defined time intervals include 37°C log phase (37°C Log), cold shock response (CSR), and two adapted growth phases at 18°C or 15°C (Log I and Log II), indicated by the red arrows. C.S.: cold shock.

*B. subtilis* was analyzed after shift to 18°C, the lowest temperature permitting long-term growth (Fig. 1D), across four phases: exponential growth at 37°C, an early CSR phase (1^st^ doubling post-shift), a late CSR phase (2^nd^–6^th^ doublings), and an adapted phase at 18°C (7^th^–11^th^ doublings) (Fig. 2B). *E. coli* was analyzed both after shift to 18°C, for direct comparison to *B. subtilis,* and 15°C, where the CSR was expected to be more pronounced. The CSR period was defined as the 1^st^ doubling post-shift, followed by the next 10 doublings as the adapted phase (Fig. 2C-D). Gene fitness values across these stages for both species are summarized in Tables S2-S6 (see Fig. S5 and Methods).

### *B. subtilis* early-stage CSR: Cell envelope remodeling and DNA damage repair

We identified 65 genes with significant fitness defects (fitness < 0.4, adjusted p < 0.05; Table S3C) and high consistency between their two biological replicates during the early stage of CSR in *B. subtilis* (Fig. 3A). Of these, 32 genes exhibited negative fitness values (i.e., mutants were depleted more than expected from growth arrest alone), suggesting potential cell lysis. Gene Ontology (GO) analysis revealed functional enrichment in teichoic acid metabolism (e.g., *ltaSB, dltA-E*), flagellar function (e.g., *hag, flhA*, *fliS*, *flhP*, *sigD*), branched-chain amino acid metabolism (e.g., *bkd* genes), and DNA recombination and repair (e.g., *recA*, *recF*, *recO*, *recG*, *ruvA*) (Fig. 3B). These results highlight two critical processes during early CSR in *B. subtilis*: remodeling cell envelope (including cell wall and membrane) and preserving genome stability.

**Figure 3.**
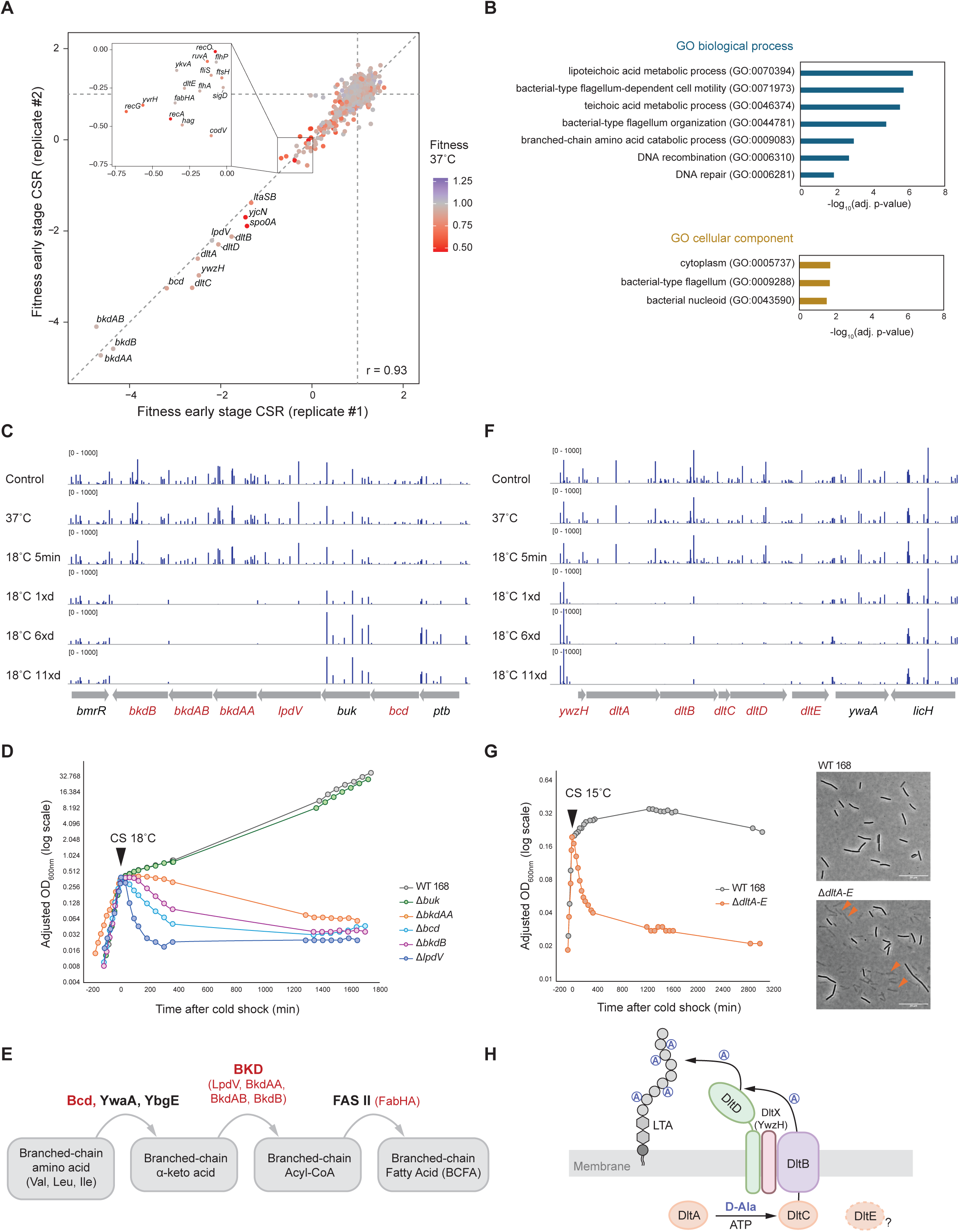
Early stage of the cold shock response (CSR) in *B. subtilis*. **(A)** Scatter plot comparing gene-specific fitness values from two biological replicates during the early stage of CSR (1^st^ doubling after cold shock). Genes with strongly decreased fitness are labeled. Colors indicate gene fitness during exponential growth at 37°C. r: Pearson’s correlation coefficient; grey dashed lines: *x* = 1, *y* = 1, and *y = x*. **(B)** Gene Ontology (GO) analysis of the 32 genes with the lowest fitness values during early CSR (fitness < 0, *p* < 0.05), showing the most significantly enriched biological processes and cellular components (see Methods). *Adj p-values*: adjusted p-values from Fisher’s exact test with false discovery rate correction. **(C)** Normalized Tn-seq signals mapped to the *bkd* operon and surrounding regions, averaged from two biological replicates. Rows represent samples collected at *t_0_* (control), 37°C before cold shock, and at 5 min, 1, 6, or 11 doublings after cold shock to 18°C. Genes with strong fitness defects are highlighted in red. **(D)** Growth curves of *B. subtilis* 168, and various single-gene deletion mutants of the *bkd* operon before and after cold shock. CS 18°C: cold shock to 18°C. **(E)** *B. subtilis* branched-chain fatty acid (BCFA) biosynthesis pathway. Branched-chain amino acids are converted to α-keto acids by branched-chain amino acid aminotransferases (BCATs) YwaA or YbgE, or alternatively by the NAD⁺-dependent dehydrogenase Bcd. The BKD complex then converts these intermediates to branched-chain acyl-CoA esters, precursors for BCFA synthesis via the FAS II pathway (72). Enzymes whose mutants showed strong fitness defects after cold shock are highlighted in red. Among them, FabH initiates FAS II; of the two isoenzymes in *B. subtilis* (FabHA and FabHB), only FabHA mutant showed cold sensitivity (Table S3). **(F)** Normalized Tn-seq signals mapped to the *dlt* operon and nearby genes, averaged from two biological replicates. **(G)** Left: Growth curves of *B. subtilis* 168 and the Δ*dltA-E* mutant before and after cold shock. CS 15°C: cold shock to 15°C. Right: Microscope images of wild-type and Δ*dltA-E* cells at 290 and 240 min after cold shock, respectively. Arrows highlight dead cells in the mutant. **(H)** Schematic of the *B. subtilis* lipid teichoic acid (LTA) D-alanylation pathway, adapted from (33).

A major cluster of hits mapped to the *bkd* operon (*ptb*-*bcd*-*buk*-*lpdV*-*bkdAA*-*bkdAB*-*bkdB*; Fig. 3C), whose expression is induced upon temperature downshift (25). Its transcriptional activator, *bkdR*, has previously been linked to cold adaptation (26). Most genes in this operon are involved in the branched-chain amino acid (BCAA) degradation pathway (27). BCAAs provide key precursors for branched-chain fatty acid (BCFA) synthesis, which Gram-positive bacteria use to maintain membrane fluidity (28, 29). To test its role in CSR, we constructed deletion mutants of representative genes. While all mutants grew normally at 37°C, most (except *buk*) underwent pronounced lysis upon cold shock to 18°C (Fig. 3D), consistent with Tn-seq data (Fig. 3C). Among them, *bcd* encodes a dehydrogenase that converts BCAAs to α-keto acids (Fig. 3E). Although the aminotransferases *ywaA* and *ybgE* can perform similar functions, their mutants showed much milder or no cold-induced defects (Table S3). LpdV, BkdAA, BkdAB and BkdB form the branched-chain keto acid dehydrogenase (BKD) complex, which converts α-keto acids to branched-chain acyl-CoA esters, the direct precursors of BCFAs (Fig. 3E). Together, these results indicate that remodeling lipid composition via BCFA synthesis is essential for maintaining membrane fluidity during CSR to counteract the rigidifying effects of low temperature, supporting earlier evidence for the importance of isoleucine in membrane adaptation to cold (28).

Another strong hit was the D-alanylation pathway of teichoic acids (DLT) (Fig. 3F). The *B. subtilis dlt* operon (*ywzH-dltA-dltB-dltC-dltD-dltE*) mediates D-alanine esterification of both lipoteichoic acid (LTA) and wall teichoic acid (WTA) (30–33). Tn-seq revealed strong depletions of *dlt* mutants shortly after cold shock (Fig. 3F). To validate this, we constructed mutants lacking one or more *dlt* genes (Δ*dltA-E*). While the *dlt* operon was dispensable for growth at 37°C, the mutants exhibited strong lysis upon cold shock to 15°C (Fig. 3G). In contrast, they showed no significant lysis after shifting to 18°C (Fig. S1B), despite negative fitness values from Tn-seq (Fig. 3A). We attribute this discrepancy to potential temperature fluctuations during the Tn-seq experiments, suggesting that *dlt* mutants are highly sensitive to small temperature changes within this range. Previous studies have revealed pleiotropic roles of teichoic acid D-alanylation (Fig. 3H) in regulating autolysis (34), resistance to cationic antibiotics (35, 36), and virulence in Gram-positive pathogens (37, 38). However, the mechanism underlying the cold sensitivity of *dlt* mutants remains unclear.

Key DNA recombination and repair factors were also critical during the early stage of CSR, including *recA*, *recF*, *recO*, *recG*, and *ruvA* (Fig. 3A). In *B. subtilis*, RecA plays a central role in DNA double-strand break repair: the AddAB complex first generates single-stranded DNA (ssDNA), RecFOR then loads RecA onto ssDNA to form a D-loop, and RecG or RuvAB promotes D-loop migration and strand extension (39). The reliance on these pathways likely reflects accumulation of ssDNA or double-stranded ends at collapsed replication forks. Consistently, Tn-seq-derived DNA dosage profiles revealed strong inhibition of DNA replication after cold shock (Fig. S2). During exponential growth at 37°C, the *ori/ter* ratio (i.e., the ratio of DNA dosage near the origin of replication to those near the terminus) was ∼4 (Fig. S2B), indicative of active multi-fork replication. Six hours after cold shock, the profile showed a plateau near the *oriC* followed by a gradual decrease toward the terminus (Fig. S2D), consistent with widespread replication fork pausing. After adaptation to 18°C, the dosage profile recovered to a V-shape (Fig. S2E-F) but with a lower *ori*/*ter* ratio, as expected due to reduced growth rate. The transient disruption of replication progression after cold shock may necessitate recombination- and repair-mediated fork restart during early adaptation.

### *B. subtilis* late-stage CSR: Post-transcriptional regulation

During the late stage of CSR (2^nd^–6^th^ doubling after shift to 18°C), we identified 106 genes with significant fitness defects (fitness < 0.6, adjusted *p* < 0.05) (Fig. 4A; Table S3D). GO enrichment analysis revealed key functional categories, including nucleic acid metabolism (e.g., *rsmA, cshB, yfmL*, *rimF, rnr, rluB*), cell division (e.g., *ezrA, minJ, divIVA, minC, ftsE*), ribosome biogenesis and translation (e.g., *rlbA, bipA, obgE*, *efp, prmC*), and DNA repair (e.g., *recA*, *recFOR, ruvAB, addA, recG*) (Fig. 4B). These top hits were highly enriched for RNA binding and catalytic activity, and strongly associated with ribonucleoprotein complexes, ribosomes, and nucleoid (Fig. 4B). The growing involvement of post-transcriptional processes suggests a shift toward reprogramming gene expression during late-stage CSR.

**Figure 4.**
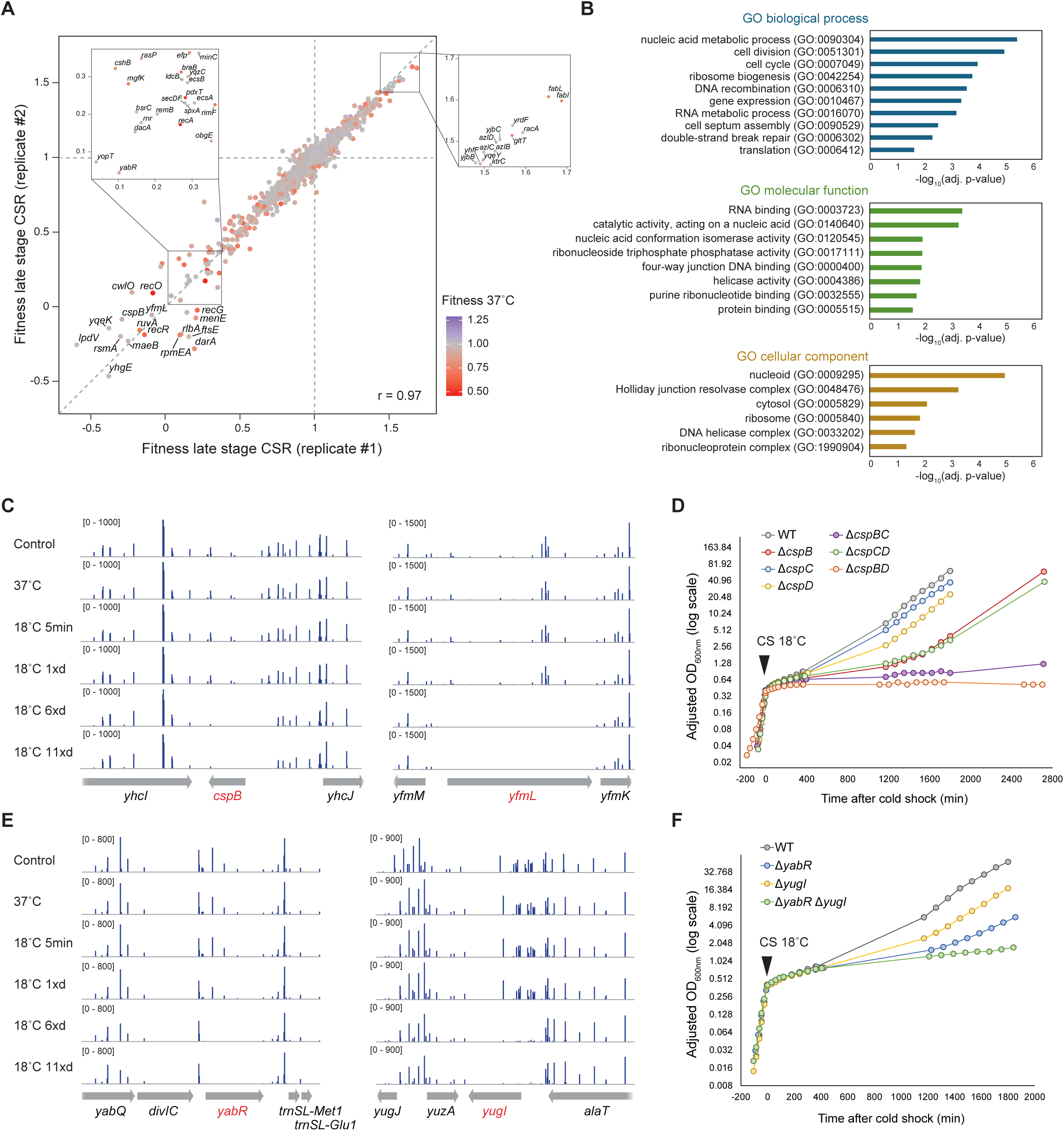
Late stage of the cold shock response (CSR) in *B. subtilis*. **(A)** Scatter plot comparing gene-specific fitness values from two biological replicates during the late stage of CSR (5 doublings after the early stage). Genes with strongly decreased or increased fitness values are labeled. Colors indicate gene fitness during exponential growth at 37°C. r: Pearson’s correlation coefficient; grey dashed lines: *x* = 1, *y* = 1, and *y = x*. **(B)** Gene Ontology (GO) analysis of 106 genes with the lowest fitness values during late CSR (fitness < 0.6, *p* < 0.05), showing the most significantly enriched biological processes, molecular functions, and cellular components (see Methods). *Adj p-values*: p-values from Fisher’s exact test adjusted with false discovery rate correction. **(C)** Normalized Tn-seq signals mapped to *cspB* (left) or *yfmL* (right) with neighboring genes, averaged from two replicates. Rows represent samples collected at t_0_ (control), 37°C before cold shock, and at 5 min, 1, 6, or 11 doublings after cold shock to 18°C. **(D)** Growth curves of *B. subtilis* 168 (WT) and single or double *csp* deletion mutants before and after cold shock. CS 18°C: cold shock to 18°C. Slow-growing strains were monitored for a longer duration than WT, Δ*cspC* and Δ*cspD*. **(E)** Normalized Tn-seq signals mapped to *yabR* (left) or *yugI* (right) with neighboring genes, averaged from two replicates. **(F)** Growth curves of *B. subtilis* 168 (WT) and single or double Δ*yabR* and Δ*yugI* mutants before and after cold shock. CS 18°C: cold shock to 18°C.

RNA-binding proteins (RBPs) formed a major group of late-stage CSR factors, reflecting the temperature sensitivity of RNA structure, metabolism and RNA-protein interactions. Among them, the cold shock proteins (Csps) and DEAD-box RNA helicases were key ones (Fig. 4C-D). Unlike *E. coli*, where Csps are dispensable at 37°C (6), *B. subtilis* requires at least one of its three paralogs (*cspB*, *cspC*, and *cspD*) for viability (40). Growth analysis of *csp* mutants revealed a functional hierarchy upon cold shock, with *cspB* being most critical, followed by *cspD* and *cspC* (Fig. 4D), consistent with prior work at 15°C (40). The Δ*cspB* mutant showed delayed adaptation, while Δ*cspBD* and Δ*cspBC* double mutants failed to grow at 18°C within 2 days (Fig. 4D). Similarly, two of the four DEAD-box helicases, CshB and YfmL, were strongly required during late-stage CSR (Fig. 4C, S1C), as previously reported at 16°C (41). The major helicase, CshA, which facilitates RNA decay in degradosome (42), showed strong depletion in Tn-seq even at 37°C (Table S2), suggesting a critical role under normal conditions. In contrast, disruption of *deaD* had no detectable effect throughout CSR (Table S3). Thus, Csps and DEAD-box helicases act as central regulators of RNA metabolism during cold adaptation, with paralogs providing both overlapping and distinct functions.

Additional factors involved in ribosomal function and translational control were also important during late-stage CSR. The large ribosomal subunit protein bL31A (*rpmEA*) and bL32 (*rpmF*), although dispensable at 37°C, showed significant fitness defects after cold shock (Fig. S1D, Table S3). Similarly, mutants lacking the translation termination regulator PrmC (*prmC*) or elongation factor EF-P (*efp*) exhibited delayed adaptation during late-stage CSR, with little growth defect at 37°C (Fig. S1E). These cold-specific phenotypes indicate that temperature downshift perturbs multiple stages of protein synthesis, spanning ribosome biogenesis, elongation, and termination, and renders these factors essential during CSR.

Interestingly, two uncharacterized S1 domain-containing proteins, YabR and YugI, also exhibited significantly lower fitness following cold shock (Fig. 4E), and the double mutant displayed a synergistic defect (Fig. 4F), suggesting partial functional redundancy. The conserved S1 domain in YabR and YugI implies RNA-binding activity, and recent studies in *B. subtilis* suggest they may interact with the 30S ribosomal subunit (43), indicating roles in ribosome biogenesis and/or translational regulation. Although it is unclear whether their ribosome association is temperature-dependent, modulating these auxiliary factors may reshape the translational landscape by altering the ribosome itself upon cold shock.

### *B. subtilis* 18°C continuous growth: fewer critical genes identified

During the subsequent five doublings, wild-type *B. subtilis* adapted to low temperature and resumed exponential growth at 18°C (Fig. 2B). Fitness analysis identified 27 genes with significant fitness defect (fitness < 0.7, adjusted p < 0.05) during adapted growth (Fig. 5A), substantially fewer than during the CSR. This is partially due to the depletion of some CSR-essential mutants before reaching adaptation (Tables S2, S3), resulting in underestimation of critical factors for continuous growth. Nevertheless, a Multidimensional Scaling (MDS) plot, which visualizes global dissimilarities across samples, revealed a stronger shift in Tn-seq profiles during the late-stage CSR than during extended growth at 18°C (Fig. 5B; see Methods). Consistently, gene-specific fitness values showed greater variability during CSR than during continuous growth at 37°C or 18°C (Fig. 5C-E). Together, these results indicate that a broader set of factors is required for initial cold shock adaptation than for long-term growth at low temperature, highlighting the importance of cold-shock-specific mechanisms during the ∼16-hour adaptation period.

**Figure 5.**
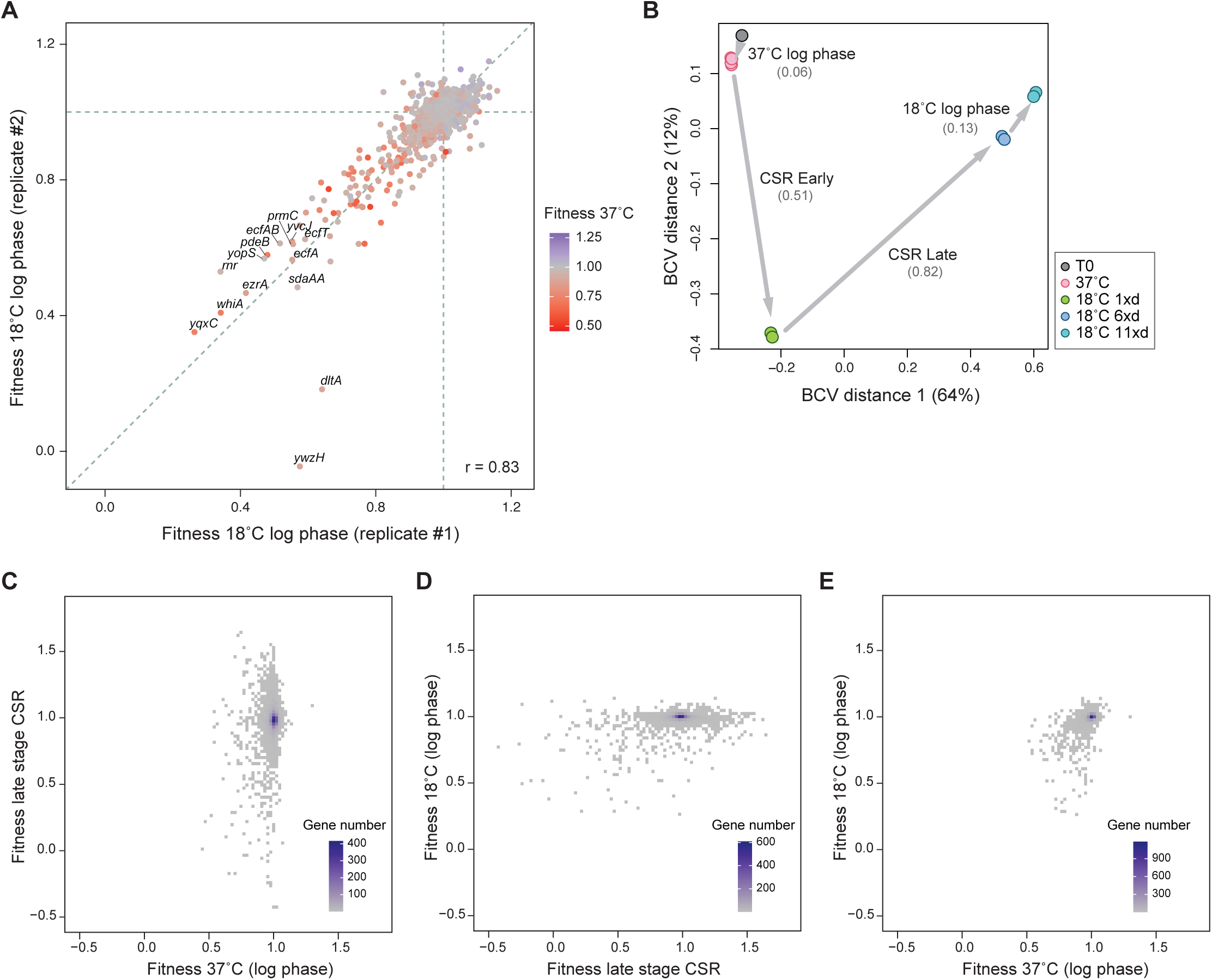
Continuous growth at 18°C after CSR in *B. subtilis*. **(B)** Scatter plot comparing gene-specific fitness values from two biological replicates during adapted exponential growth at 18°C (5 doublings after the late stage of CSR). Genes with strongly decreased fitness are labeled. Colors indicate fitness values during exponential growth at 37°C. r: Pearson’s correlation coefficient; grey dashed lines: *x* = 1, *y* = 1, and *y = x*. **(C)** Multidimensional scaling (MDS) plot of *B. subtilis* Tn-seq profiles. MDS plots visualize sample dissimilarities, conceptually similar to Principal Component Analysis (PCA) but based on pairwise distances. Samples (with biological replicates) were collected at t_0_ (control), 37°C before cold shock, 5 minutes after cold shock, and after 1, 6, or 11 doublings at 18°C. The 5 min post-shift samples are grouped with the 37°C samples, as they are not statistically distinct (Fig. S5F). Dimension 1 (BCV distance 1) accounts for 64% of the variance in mutant abundance, and Dimension 2 (BCV distance 2) accounts for 12%. Grey arrows mark critical time intervals: 37°C log phase, early and late stages of cold shock response (CSR), and the adapted log phase at 18°C. Numbers below labels indicate the average pairwise Euclidean distances, reflecting the level of dissimilarity between groups. **(C-E)** Scatter plots comparing gene-specific fitness values (averaged from 3 biological replicates) between selected time intervals, based on Tn-seq analysis. Colors represent gene density.

Persistent growth defects at 18°C were enriched in mutants disrupting cell envelope (e.g., *bkd* and *dlt* operons) and translation or ribosome regulation (e.g., *rsmA, nusG, bipA, rluB, rnr, prmC*) (Table S3E). These mutants likely fail to fully resume growth (e.g., due to cell lysis), exhibit markedly delayed recovery, or experience a continuous growth defect even after adaptation.

### *E. coli* vs. *B. subtilis*: Broadly conserved and species-specific CSR factors

Compared to *B. subtilis*, fewer genes were found essential for CSR (1^st^ doubling post-shock) in *E. coli*. After shifting to 18°C, 19 genes showed significantly reduced fitness during CSR (fitness < 0.6, adjusted p < 0.05), and 39 genes during sustained growth at 18°C (fitness < 0.85, adjusted p < 0.05; Tables S5). Cold shock to 15°C yielded 6 and 38 such genes, respectively (Tables S6). About 70% of these genes overlapped between two cold shock conditions (Fig. 6A), indicating a shared core cold adaptation program. MDS plots of Tn-seq profiles revealed smaller phenotypic shifts during CSR than during adapted growth at 18°C or 15°C (Fig. 6B-C). The overall fewer and milder mutant phenotypes in *E. coli* than *B. subtilis* suggest that a smaller gene set is required for the acute CSR in *E. coli*.

**Figure 6.**
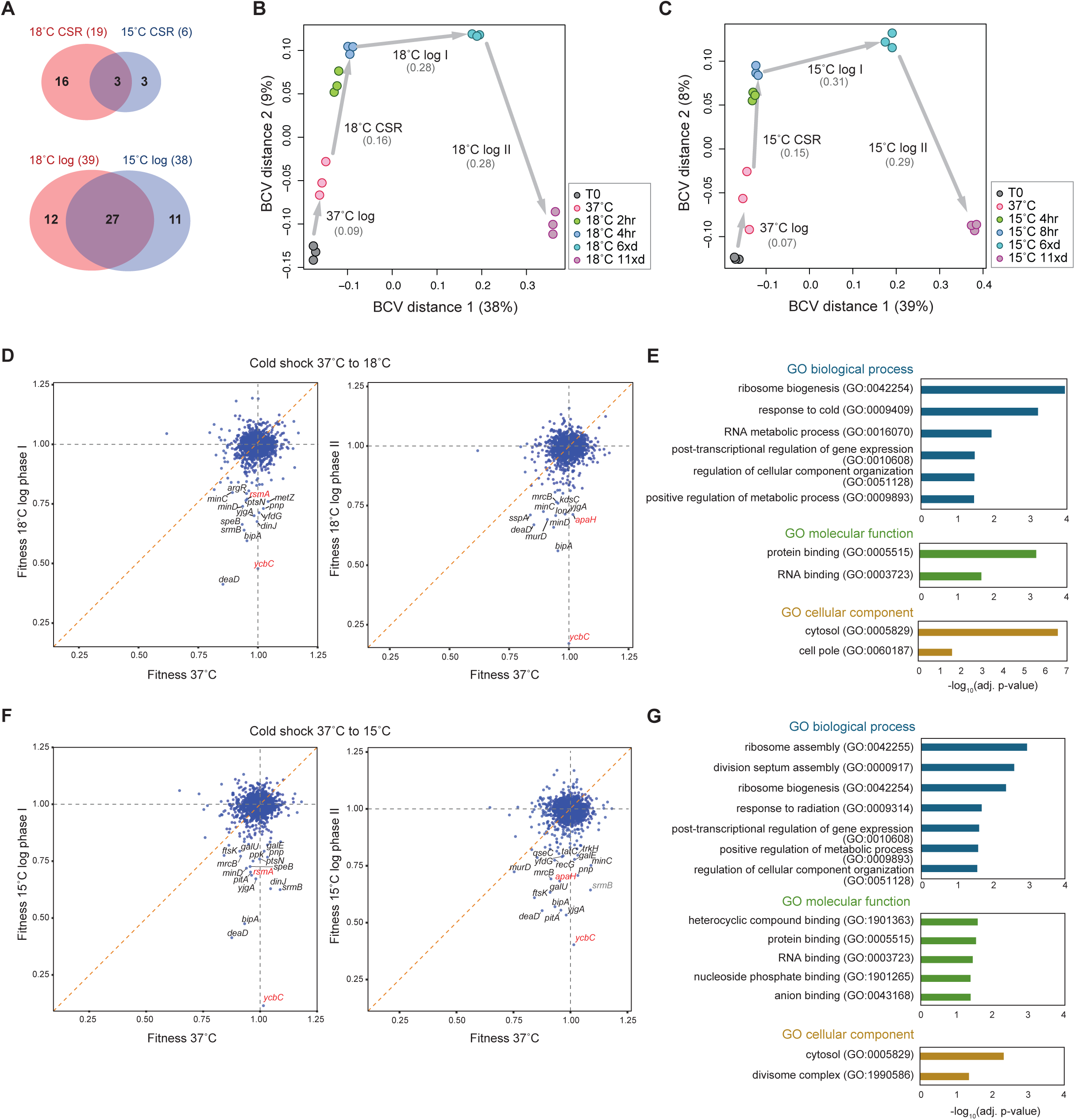
RB-Tnseq profiles of *E. coli* cold shock response (CSR) **(A)** Venn diagrams comparing genes important for *E. coli* CSR upon shift from 37°C to 18°C versus 37°C to 15°C (top), and during exponential growth at 18°C versus 15°C after adaptation (bottom). Numbers indicate gene counts in each group. **(B)** Multidimensional scaling (MDS) plot of *E. coli* RB-Tnseq profiles before and after cold shock from 37°C to 18°C. Samples were collected at t_0_ (control), 37°C before cold shock, and at 2 hours, 4 hours (1 doubling) and 6 or 11 doublings after cold shock (3 biological replicates). Dimension 1 (BCV distance 1) accounts for 38% of the variance in mutant abundance, and Dimension 2 (BCV distance 2) accounts for 9%. Grey arrows mark key time intervals: 37°C log phase, CSR, and two adapted growth phases at 18°C. Numbers below labels indicate the average pairwise Euclidean distances, reflecting the level of dissimilarity between groups. **(C)** MDS plots of *E. coli* RB-Tnseq profiles before and after cold shock from 37°C to 15°C. Samples were collected at t_0_ (control), 37°C before cold shock, and at 4 hours, 8 hours (1 doubling) and 6 or 11 doublings after cold shock. Dimension 1 and 2 explain 39% and 8% of the total variance in mutant abundance, respectively. Grey arrows mark 37°C log phase, CSR, and two adapted growth phases at 15°C. **(D)** Scatter plots comparing gene-specific fitness values (averaged from 3 biological replicates) during exponential growth at 37°C vs. adapted growth phases at 18°C. Genes with strong fitness defects at 18°C are labeled. Highlighted genes are discussed in the main text. Grey dashed lines: *x* = 1, *y* = 1; orange dashed line: *y = x*. **(E)** Gene Ontology (GO) analysis of 48 genes important during CSR (fitness < 0.6, p < 0.05) or continuous growth at 18°C (fitness < 0.85, p < 0.05), showing the most significantly enriched biological processes, molecular functions, and cellular components (see Methods). **(F)** Scatter plots comparing gene-specific fitness values (averaged from 3 biological replicates) during exponential growth at 37°C vs. adapted growth phases at 15°C. Genes with strong fitness defects at 15°C are labeled. Grey dashed lines: *x* = 1, *y* = 1; orange dashed line: *y = x*. **(G)** GO analysis of 41 genes important during CSR (fitness < 0.6, p < 0.05) or continuous growth at 15°C (fitness < 0.85, p < 0.05).

Tn-seq is limited in capturing phenotypes during the brief, nongrowing acclimation phase of *E. coli* CSR. Therefore, we combined key factors identified from both CSR and continuous cold growth for further analysis. GO analysis of these genes revealed key functional categories, including RNA metabolism (e.g., *srmB, deaD*, *nusA*, *pnp*, *apaH, rsmA* and *rsmH*), ribosome biogenesis (e.g., *lepA*, *bipA*, *yjgA*), and cell division (e.g., *minC* and *minD*) (Fig. 6D-G). Similar to *B. subtilis*, post-transcriptional regulation, particularly of RNA metabolism and translation, play a central role in both CSR and continuous cold growth. Conserved factors in both species include DEAD-box RNA helicases (SrmB, DeaD in *E. coli*, and CshB, YfmL in *B. subtilis*), rRNA methyltransferases (RsmA, RsmH), and ribosome assembly proteins (BipA). Interestingly, some CSR factors perform conserved functions despite limited sequence similarity, suggesting evolutionary divergence (see Discussion).

We also identified *E. coli*-specific CSR factors. Among the strongest hits was *ycbC* (*elyC*), which is critical for cell wall and surface polysaccharide biogenesis (44). *elyC* mutants showed significantly reduced fitness after cold shock to both 18°C and 15°C (Fig. 6D,F), consistent with the previous observations of impaired growth and increased lysis at room temperature (44). Moreover, *mrcB*, encoding Penicillin-binding protein 1B (PBP1B), was reported as synthetically lethal with *elyC* deletion at room temperature (44), and *mrcB* transposon mutants also exhibited reduced fitness after cold shock in our Tn-seq results (Fig. 6D,F), suggesting their functional connection in remodeling cell envelope during cold adaptation.

### Conserved roles of rRNA methyltransferases in cold adaptation

Ribosomal RNA (rRNA) modifications are critical for ribosome biogenesis and function. Our Tn-seq analysis identified two 16S rRNA methyltransferases, RsmA and RsmH, as key factors in cold adaptation of both species. RsmA (or KsgA) di-methylates A1518 and A1519 in *E. coli* (45), and RsmH methylates C1402 at the ribosomal P site (46). These modifications are broadly conserved across all domains of life (47).

To test their functions under cold stress, we constructed single and double deletion mutants of *rsmA* and *rsmH*. In *B. subtilis*, the Δ*rsmA* mutant showed a pronounced delay in cold adaptation and slower growth at 18°C (Fig. 7A, S7A), whereas the Δ*rsmH* mutant displayed milder phenotypes (Fig. 7A, S7A). Strikingly, the double mutant exhibited a synergistic defect, failing to recover even after 4 days at 18°C (Fig. 7A, S1F). Its slower growth rate at 37°C relative to wild type suggests housekeeping roles of these modifications under non-stress conditions. This synergy is conserved in *E. coli*, where the Δ*rsmA* Δ*rsmH* double mutant showed a longer acclimation phase and sustained growth defect at both 15°C (Fig. 7B) and 18°C (Fig. S7B). In contrast, the single mutants had much weaker phenotypes in *E. coli* than in *B. subtilis* (Fig. 7A-B, S7A-B).

**Figure 7.**
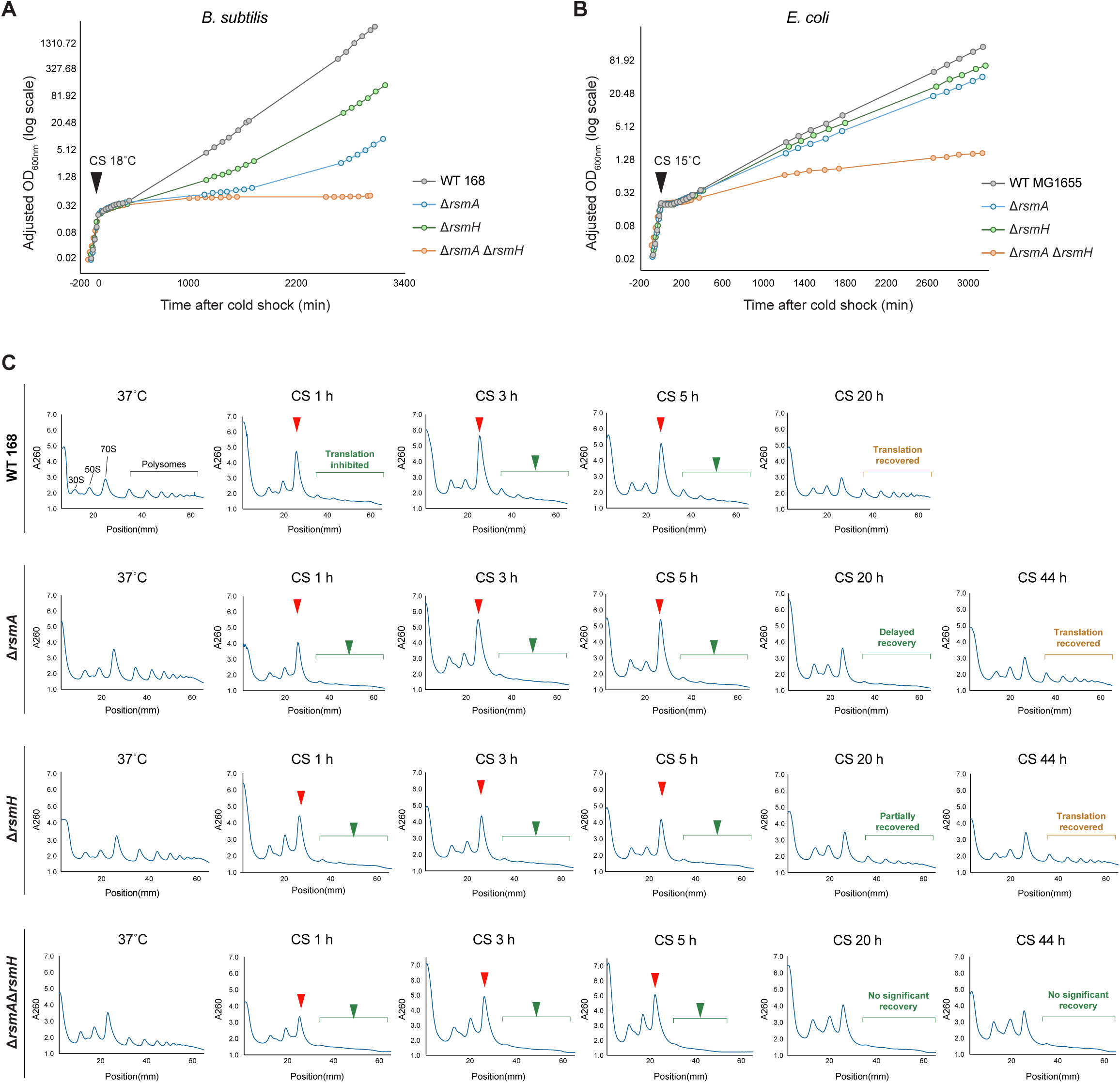
Roles of RsmA and RsmH in cold shock adaptation of *B. subtilis*. **(A-B)** Growth curves of *B. subtilis* 168 (WT), Δ*rsmA,* Δ*rsmH*, and the Δ*rsmA* Δ*rsmH* double mutant **(A)**, or *E. coli* MG1655 (WT) and corresponding single and double *rsm* mutants **(B)**, before and after cold shock. CS 18°C: cold shock to 18°C; CS 15°C: cold shock to 15°C. **(D)** Polysome profiles of *B. subtilis* WT, Δ*rsmA*, Δ*rsmH,* and Δ*rsmA* Δ*rsmH* mutants at selected time points before and after cold shock. X-axis: position from the top of the centrifuge tube; Y-axis: UV absorbance at 260nm (A260). In the WT 37°C profile, peaks corresponding to 30S and 50S ribosomal subunits, 70S monosomes, and polysomes are indicated. Cold shock causes an accumulation of 70S monosomes (red arrows) and depletion of polysomes (green arrows). In WT cells, translation recovered after ∼20 hours, whereas Δ*rsmA* and Δ*rsmH* single mutants showed delayed recovery. The double mutant failed to restore translation even after 44 hours.

The most likely explanation for the role of these rRNA methyltransferases is their impact on ribosome assembly, which is intrinsically cold sensitive (48), as previously shown for RsmA in *E. coli* (49). To test this, we analyzed polysome profiles of wild-type and mutant strains using sucrose gradient sedimentation (Figs. 7C, S7C). In wild-type cells of both species, cold shock caused an accumulation of 70S monosomes and depletion of polysomes (translating ribosomes), indicating global translation inhibition. The increase in 70S ribosomes may reflect stalling at initiation sites or accumulation of inactive particles. These changes peaked at 3 hours after shifting to 18°C in *B. subtilis* and at 10–30 minutes in *E. coli*, and profiles then recovered as cells adapted to 18°C (Figs. 7C, S7C). In *B. subtilis*, *rsm* single mutants showed delayed polysome recovery, with Δ*rsmA* having a stronger defect (Fig. 7C). The double mutant failed to restore translation even after 2 days (44 hours; Fig. 7C), consistent with its severe growth defect (Fig. 7A). As expected, *E. coli rsm* single mutants did not show major differences compared to wild type, in line with their weak phenotypes (Fig. S7C). Importantly, we did not observe significant enrichment of additional peaks indicative of severe ribosome assembly defects in any sample (48) (Figs. 7C, S7C).

Together, these results suggest that RsmA and RsmH contribute to global translation recovery during CSR, likely by promoting 70S ribosome maturation and/or enabling efficient translation at low temperatures. Although no aberrant assembly intermediates were detected, these modifications may act as quality-control checkpoints, preventing defective ribosomes from engaging in translation (50). Moreover, the modifications may also influence downstream processes such as translation fidelity. Future genome-wide translation profiling will be required to test these possibilities.

## Discussion

The cold shock response (CSR) involves multiple cellular processes that act on thermosensitive features such as membrane fluidity, nucleic acid structure, protein folding, and ribosome assembly and function. They serve as both triggers and targets of CSR, making it challenging to disentangle the functional contribution of each pathway. Although previous studies have identified many cold-inducible proteins (CIPs) whose expression increase after cold shock, their induction does not necessarily imply functional importance.

Our Tn-seq-based phenotypic profiling provides the first systematic functional analysis across the full temporal progression of CSR. In contrast to rapid stress responses such as the heat shock response, which activates within minutes, CSR can unfold over hours. In *B. subtilis*, where CSR spans ∼16 hours (Fig. 2B), our time-resolved approach uncovered stage-specific gene requirements. During the early phase of CSR, genes involved in maintaining membrane fluidity and cell wall integrity are critical (Fig. 3), highlighting the cell envelope as the first line of defense against environmental stress. In the late phase, adaptation is predominantly mediated by post-transcriptional mechanisms (Fig. 4), reflecting the need to reprogram gene expression to resume growth. Cold stress broadly impairs transcription and translation, with translation emerging as the major bottleneck of gene expression. Thus, remodeling post-transcriptional regulation become critical to optimize the limited protein synthesis capacity. Additionally, genes involved in DNA recombination and repair are required throughout CSR (Fig. 3, 4), suggesting persistent replication challenges under cold stress. Overall, this gene function profile provides a comprehensive view of the coordinated cellular processes that underlie CSR over time.

While our primary focus was on identifying positive CSR factors (i.e., genes required to maintain fitness), we also uncovered putative negative factors, whose disruption enhanced growth during cold adaptation. A notable example in *B. subtilis* involves the paralogous enoyl-acyl carrier protein (ACP) reductases FabI and FabL (Fig. S6A-B), which catalyze the final step of fatty acid elongation cycle (51). Mutants in either gene exhibited significantly elevated fitness (Fig. 4A; adjusted p< 0.05) under cold shock. Partial loss of FabI/L function may alter fatty acid composition, shifting toward shorter or branched-chain fatty acids, thereby increasing membrane fluidity. Such changes could transiently benefit cells coping with cold-induced membrane rigidification.

CSR strategies vary across species. One notable difference lies in their post-shock growth behaviors. *E. coli* exhibits a no-growth lag phase immediately following cold shock, a pattern also observed in *Vibrio cholerae*, *Listeria monocytogenes*, and *Mycobacterium smegmatis* (52–54). In contrast, *B. subtilis* undergoes a gradual decline in growth rate before slowly recovering to a new steady state, a pattern shared by *Lactococcus lactis* and *Enterococcus faecalis* (55, 56). The presence or absence of a lag phase may reflect differences in the extent or mechanism of translation inhibition. The molecular “brake” that governs this acclimation phase remains unidentified, but may involve intrinsic ribosome sensitivity or specialized regulatory factors. This regulatory brake also partially contributes to the observed differences in CSR dynamics, as *E. coli* resumes full-speed growth at low temperatures more rapidly than *B. subtilis*. Importantly, these phenotypic variations across species cannot be fully explained by their optimal growth temperatures or phylogenetic relationships.

Direct comparison of *E. coli* and *B. subtilis* phenotypic profiles revealed both conserved and species-specific CSR components. A large group of conserved factors are involved in post-transcriptional control. In both species, RNA binding proteins (RBPs) play central roles in CSR, including the broadly conserved cold shock proteins, RNA helicases (e.g., DEAD-box helicases), RNA modification enzymes (e.g., rRNA methyltransferases RsmA, RsmH), and ribosome assembly factors (e.g., BipA). These findings highlight ribosomes and RNAs as key sensors and effectors in CSR, and underscore the evolutionary conservation of RNA remodeling and translational reprogramming as core strategies for cold adaptation.

In some cases, functionally conserved CSR factors share little sequence similarity across species. For example, the *E. coli* gene *apaH*, which encodes a diadenosine tetraphosphatase, is one of the strongest CSR hits (Fig. 6D, 6F, S6C). ApaH hydrolyzes diadenosine tetraphosphate (Ap_4_A), a side product of aminoacyl-tRNA synthetases (57, 58). Ap_4_A has been proposed as a stress-related alarmone, with intracellular levels increasing upon oxidative stress, heat shock and antibiotic exposure (59–61), although its physiological role remains unclear. In addition to hydrolyzing Ap_4_A, ApaH can decap RNAs bearing Ap_4_ or Gp_4_ at their 5’ ends (62), suggesting a broader role in RNA metabolism. In *B. subtilis*, the HD-domain enzyme YqeK has been shown to degrade Ap_4_A (63), but no evidence currently supports a role in RNA decapping. YqeK homologs are restricted to Gram-positive bacteria lacking ApaH homologs (64). Despite their divergent domain structures and evolutionary origins, both *apaH* and *yqeK* mutants exhibited CSR-specific fitness defects (Fig. S6C-D), suggesting a conserved role in Ap_4_A turnover and/or RNA decapping to regulate RNA stability under cold stress. Further studies are needed to clarify their molecular functions and contributions to CSR.

Specifies-specific CSR strategies often target the cell envelope, reflecting fundamental differences between Gram-positive and Gram-negative bacteria. For example, the *bkd* pathway for branched-chain fatty acid biosynthesis and the *dlt* pathway for D-alanylation of teichoic acids (Fig. 3E,H) are conserved in Gram-positive species, but absent in *E. coli*, which relies on alternative mechanisms. Thus, while the molecular strategies diverge, both groups remodel their envelopes to restore membrane fluidity and maintain cell wall integrity during cold adaptation.

While our study provides new system-level insights into CSR, important gaps remain. First, essential and quasi-essential genes were excluded from our transposon screen due to low or absent insertion frequency, limiting genome-wide coverage. CRISPR interference (CRISPRi)-based approaches could overcome this limitation by enabling knockdown of essential genes. However, the impact of temperature on CRISPRi efficacy needs to be assessed. Second, gene redundancy, such as among the cross-regulated RNA-binding proteins with conserved domains, can mask phenotypes in single-gene mutants. Finally, our understanding of CSR in non-model organisms remain limited, particularly in species with distinct temperature tolerances and preferences, such as extremophiles. Notably, cold shock proteins were found to be critical for virulence in the foodborne pathogen *Listeria monocytogenesis* (65), which can grow at refrigeration temperatures. Expanding CSR studies to a broad range of bacterial species will not only uncover new regulatory mechanisms but also inform microbial engineering and development of antimicrobials.

## Materials and Methods

### Strains and growth conditions

*Bacillus subtilis* 168 and *E. coli* K-12 MG1655 were used as wild-type strains. In *B. subtilis*, single-gene deletions were constructed by transforming wild-type cells with ∼1kb flanking regions surrounding the Kanamycin resistance cassette amplified from a previously described library (66), followed by Cre/lox excision to obtain markerless mutants. In *E. coli*, single-gene deletions were generated by P1 phage transduction from the Keio collection (67), with subsequent marker removal by Flp recombinase expressed from plasmid pCP20 (68). All strains were grown in LB Lennox medium (Fisher Scientific) at 37°C with shaking at 240 rpm in water bath shakers. Low-temperature growth was performed by incubating cultures in shakers placed in a 4°C cold room set to the desired temperature. All the strains and oligos used in this study are listed in SI Appendix and Table S1.

### Cold shock growth measurement

For both *E. coli* and *B. subtilis*, overnight cultures grown at 37°C were inoculated into pre-warmed LB medium at an initial OD_600_ of 0.005. Cells were grown at 37°C until reaching exponential phase (OD_600_ ∼ 0.2 for *B. subtilis* and ∼ 0.3 for *E. coli*), then subjected to cold shock by rapidly mixing the 37°C culture with a defined volume of pre-chilled LB medium. Cultures were then immediately transferred to a shaker set to the target low temperature, and growth was monitored until adaptation was complete. To prevent entry into stationary phase during continuous growth, cultures were back diluted to OD_600_= 0.01 in temperature-equilibrated medium once OD_600_ reached ∼ 0.3.

### Bacillus subtilis and E. coli Tn-seq

The *B. subtilis* transposon library (22) and *E. coli* barcoded transposon library (23) were grown at 37°C and subjected to cold shock. Samples were collected at t_0_ (before growth), after 4 doublings at 37°C, shortly after cold shock (5 min for *B. subtilis*; 2 h for *E. coli* at 18°C or 4 h at 15°C), and after 1, 6, and 11 doublings post-shock. Cells were harvested by centrifugation at 5000 x g for 10 minutes and flash frozen in liquid nitrogen. Tn-seq sequencing libraries were constructed as previously described (22, 24, 69). Briefly, genomic DNA was extracted from each sample, and the transposon insertion sites (*B. subtilis*) or the associated DNA barcodes (*E. coli*) for all the individual mutants were PCR-amplified using primers compatible with Illumina adaptors. Products were sequenced using Illumina NovaSeq 6000. Tn-seq reads were then mapped to the respective genome, and a LOESS (Locally Estimated Scatterplot Smoothing) model was applied to correct for chromosomal position bias from bidirectional replication (70)(Fig. S2-S4). Gene fitness was calculated as the change in mutant abundance over defined time intervals, normalized to generation number (24).

### Gene Ontology (GO) enrichment

GO enrichment analysis of genes was performed using PANTHER Overrepresentation Test (https://geneontology.org/; GO Ontology database DOI: 10.5281/zenodo.15066566 Released 2025-03-16). Overrepresentation analysis was performed to identify enriched biological process, molecular function, and cellular component with default settings.

### Multidimensional scaling (MDS) analysis

Multidimensional scaling (MDS) was used to visualize the similarity (or dissimilarity) of Tn-seq profiles across conditions. Normalized read counts of individual genes were imported into R, and MDS plots were generated using the edgeR package (71). Distances between samples were calculated using the Euclidean method, and the first two dimensions were plotted to represent the major sources of variation.

### Polysome Profiling

200 mL of culture was vacuum filtered through a 0.22 µm nitrocellulose membrane (GVS) and flash-frozen in liquid nitrogen. Cell pellet was lysed using 10 mL canisters (QIAGEN) with 500 µL frozen lysis buffer (100mM NH_4_Cl, 10mM MgCl_2_, 5mM CaCl_2_, 20mM Tris-HCl pH 8.0, 0.1% NP-40, 0.4% Triton X-100, 1mM chloramphenicol, 100 U/mL DNase I (Roche)) for 5 cycles of 3 minutes at 15 Hz with a QIAGEN Tissue Lyser III. The canisters were re-chilled in liquid nitrogen between cycles. Pulverized lysates were thawed and clarified by centrifugation at 16000 x g for 10 minutes at 4°C. 10 A_260_ units of clarified lysates were ultracentrifuged at 35,000 rpm for 2.5 hours at 4°C in an SW-41 Ti rotor (Beckman Coulter) through 10–55% sucrose gradients prepared in gradient buffer (100mM NH4Cl, 10mM MgCl2, 5mM CaCl2, 20mM Tris-HCl pH 8.0, 100 µg/mL chloramphenicol, 2mM DTT). Gradients were fractionated using a Biocomp Gradient Station (Biocomp Instruments), and polysome profiles were recorded by continuous A_260_ monitoring with a TRIAX™ Flow Cell (Biocomp Instruments).

**Further details can be found in the SI Appendix.**

## Data availability

All sequencing data from this study will be deposited in the NCBI SRA, and all custom code will be released on GitHub, prior to formal journal submission.

## Declaration of interests

The authors declare no competing interests.

## Acknowledgements

We thank Drs. Carol Gross and Alan Grossman for generously sharing strains and materials, and Drs. Byoung-Mo Koo and Jordi van Gestel for their help with transposon sequencing experiments and data analysis. We also thank Drs. Joe Sanfilippo and Raven Huang for providing access to equipment. Special thanks to Dr. Carol Gross for insightful feedback on the project and manuscript, and to the members of the Zhang lab for critical reading of the manuscript. This work was supported by start-up funds from the University of Illinois Urbana-Champaign. Y.Z. acknowledges support from Dr. Carol Gross (NIH R35 GM118061). V.K.M acknowledges the Biopreparedness Research Virtual Environment (BRaVE) Phage Foundry at Lawrence Berkeley National Laboratory supported by the U.S. Department of Energy, Office of Science, Office of Biological & Environmental Research under contract number DE-AC02-05CH11231.

## Notes

### Competing Interest Statement

The authors have declared no competing interest.

