## Supplemental Information for "Cross-species phenotypic profiling uncovers functional determinants of bacterial cold shock adaptation"


**This PDF file includes:**

Supplementary Materials and Methods

Figures S1 to S7

Legends for Tables S1 to S6

SI References

#### Supplementary Materials and Methods

##### Reagents and tools table

| Reagent/Resource | Reference or Source | Identifier or Catalog Number |
|---|---|---|
| **Experimental models** | | |
| *Bacillus subtilis* 168 | BGSC | 1A1 |
| *Bacillus subtilis* NCIB 3610 | BGSC | 3A1 |
| *Escherichia coli* MG1655 | (1) | N/A |
| *E. coli ΔrsmA::Kan*$^R$ | (2) | N/A |
| *E. coli ΔrsmH::Kan*$^R$ | (2) | N/A |
| *E. coli ΔrsmA* | This study | N/A |
| *E. coli ΔrsmH* | This study | N/A |
| *E. coli ΔrsmAΔrsmH* | This study | N/A |
| *B. subtilis ΔrsmA::Kan*$^R$ | (3) | BKK with locus tag numbers |
| *B. subtilis ΔrsmH::Kan*$^R$ | (3) | BKK with locus tag numbers |
| *B. subtilis ΔrsmA* | This study | N/A |
| *B. subtilis ΔrsmH* | This study | N/A |
| *B. subtilis ΔrsmAΔrsmH* | This study | N/A |
| *B. subtilis ΔdltA::Kan*$^R$ | (3) | BKK with locus tag numbers |
| *B. subtilis ΔdltC::Kan*$^R$ | (3) | BKK with locus tag numbers |
| *B. subtilis ΔdltD::Kan*$^R$ | (3) | BKK with locus tag numbers |
| *B. subtilis ΔdltA* | This study | N/A |
| *B. subtilis ΔdltC* | This study | N/A |
| *B. subtilis ΔdltD* | This study | N/A |
| *B. subtilis ΔdltA-E* | A gift from BM Koo | N/A |
| *B. subtilis ΔyabR::Kan*$^R$ | (3) | BKK with locus tag numbers |
| *B. subtilis ΔyugI::Kan*$^R$ | (3) | BKK with locus tag numbers |
| *B. subtilis ΔyabR* | This study | N/A |
| *B. subtilis ΔyugI* | This study | N/A |
| *B. subtilis ΔyabRΔyugI* | This study | N/A |
| *B. subtilis Δbuk::Kan*$^R$ | (3) | BKK with locus tag numbers |
| *B. subtilis Δbuk* | This study | N/A |
| *B. subtilis ΔbkdAA::Kan*$^R$ | (3) | BKK with locus tag numbers |
| *B. subtilis ΔbkdAA* | This study | N/A |
| *B. subtilis Δbcd::Kan*$^R$ | (3) | BKK with locus tag numbers |

| | | |
|---|---|---|
| *B. subtilis Δbcd* | This study | N/A |
| *B. subtilis ΔbkdB::Kan*$^R$ | (3) | BKK with locus tag numbers |
| *B. subtilis ΔbkdB* | This study | N/A |
| *B. subtilis ΔlpdV::Kan*$^R$ | (3) | BKK with locus tag numbers |
| *B. subtilis ΔlpdV* | This study | N/A |
| *B. subtilis ΔcshB::Kan*$^R$ | (3) | BKK with locus tag numbers |
| *B. subtilis ΔcshB* | This study | N/A |
| *B. subtilis ΔyfmL::Kan*$^R$ | (3) | BKK with locus tag numbers |
| *B. subtilis ΔyfmL* | This study | N/A |
| *B. subtilis ΔrpmEA::Kan*$^R$ | (3) | BKK with locus tag numbers |
| *B. subtilis ΔrpmEA* | This study | N/A |
| *B. subtilis ΔrpmF::Kan*$^R$ | (3) | BKK with locus tag numbers |
| *B. subtilis ΔrpmF* | This study | N/A |
| *B. subtilis Δefp::Kan*$^R$ | (3) | BKK with locus tag numbers |
| *B. subtilis Δefp* | This study | N/A |
| *B. subtilis ΔprmC::kan*$^R$ | (3) | BKK with locus tag numbers |
| *B. subtilis ΔcspB::kan*$^R$ | (3) | BKK with locus tag numbers |
| *B. subtilis ΔcspC::kan*$^R$ | (3) | BKK with locus tag numbers |
| *B. subtilis ΔcspD::kan*$^R$ | (3) | BKK with locus tag numbers |
| *B. subtilis ΔcspB* | This study | N/A |
| *B. subtilis ΔcspC* | This study | N/A |
| *B. subtilis ΔcspD* | This study | N/A |
| *B. subtilis ΔcspB ΔcspC* | This study | N/A |
| *B. subtilis ΔcspC ΔcspD* | This study | N/A |
| *B. subtilis ΔcspB ΔcspD::Kan*$^R$ | This study | N/A |
| *B. subtilis ΔcspD ΔcspB::Kan*$^R$ | This study | N/A |
| **Recombinant DNA** | | |
| pCP20 | (4) | N/A |
| pDR244 | (3) | N/A |
| **Oligonucleotides and other sequence-based reagents** | | |
| Oligos used in this study are listed in Table S1. | This study | Table S1 |
| **Chemicals, Enzymes and other reagents** | | |
| LB Broth (Powder) Lennox | Fisher Scientific | BP1427-2 |

| | | |
|---|---|---|
| GVS Nitrocellulose-Mixed Esters of Cellulose Membrane Filters | Fisher Scientific | E02WP09025 |
| DNase I recombinant, RNase-free (Roche) | Sigma-Aldrich | 4716728001 |
| Thermo Scientific™ NP-40 Surfact-Amps™ Detergent Solution | Fisher Scientific | PI85124 |
| Triton X-100 | Fisher Scientific | BP151-100 |
| 1M $MgCl_2$ | Invitrogen | AM9530G |
| 1M Tris, pH8.0, RNase free | Invitrogen | AM9855G |
| 1M $CaCl_2$ | Sigma-Aldrich | 21115-1ML |
| Ammonium chloride | Sigma-Aldrich | A9434-500G |
| DL-Dithiothreitol, DTT | Sigma-Aldrich | D9779-5G |
| Chloramphenicol | Sigma-Aldrich | C0378-25G |
| Kanamycin sulfate from Streptomyces kanamyceticus | Sigma-Aldrich | K1377-25G |
| Spectinomycin dihydrochloride pentahydrate | Sigma-Aldrich | S9007-25G |
| TissueLyser III | QIAGEN | 9003240 |
| Grinding Jar Set, S. Steel (2 x 10 mL) | QIAGEN | 69985 |
| Spectinomycin dihydrochloride pentahydrate | Sigma-Aldrich | S9007 |
| DNeasy Blood & Tissue Kit | QIAGEN | 69504 |
| QIAquick PCR Purification Kit | QIAGEN | 28104 |
| DNA Clean & Concentrator-5 | Zymo Research | D4014 |
| MmeI | New England Biolabs | R0637S |
| Quick CIP | New England Biolabs | M0525S |
| T4 DNA Ligase | New England Biolabs | M0202S |
| Q5 High-Fidelity DNA Polymerase | New England Biolabs | M0491S |
| Invitrogen™ Novex™ TBE Gels, 8% | Invitrogen | EC62152BOX |
| **Software** | | |
| Bowtie version 1.2.3 | (5) | https://bowtie-bio.sourceforge.net/index.shtml |
| IGV (Integrative Genomics Viewer) | (6) | https://igv.org/ |

| | | |
|---|---|---|
| Biostrings | Pagès H, Aboyoun P, Gentleman R, DebRoy S (2024). R package version 2.72.1 | doi:10.18129/B9.bioc.Biostrings |
| edgeR | (7) | doi:10.18129/B9.bioc.edgeR |
| GO enrichment analysis | PANTHER 19.0 | https://geneontology.org/ |
| Fiji 2.16.0 | (8) | https://imagej.net/ |

#### Strains and growth conditions

*Bacillus subtilis* 168 and *E. coli* K-12 MG1655 were used as wild-type strains. In *B. subtilis*, single-gene deletion strains were constructed by transforming wild-type cells with ~1kb flanking regions surrounding the Kanamycin resistance cassette from a previously constructed single-gene deletion library (3). To generate clean deletions, plasmid pDR244 (a temperature-sensitive plasmid constitutively expressing Cre recombinase) was introduced to excise the *lox*-flanked Kanamycin resistance cassette (3). Transformants were selected on LB agar plate supplemented with 100 µg/mL spectinomycin at 30˚C, and then streaked onto LB agar plates and incubated at 42˚C to cure the plasmid. Final clean mutants were confirmed by PCR with primers flanking the deletion site. Similarly, in *E. coli*, single-gene deletion strains were generated by P1 phage transduction of FRT-flanked deletion alleles from the Keio collection (2) to MG1655 cells. The antibiotic resistance marker was subsequently removed by expressing Flp recombinase from plasmid pCP20 (4).

All cultures were grown in LB Lennox medium (Fisher Scientific) at 37˚C with continuous shaking at 240 rpm in water bath shakers. For low-temperature growth experiments, the shakers were placed in a 4˚C cold room and set to the desired temperature (e.g., 18˚C, 15˚C, 12˚C, or 9˚C).

#### Cold shock growth measurement

All strains were cultured in LB-Lennox medium, and water-bath shakers were used to minimize temperature fluctuations. For both *E. coli* and *B. subtilis*, overnight cultures grown at 37˚C were inoculated into pre-warmed LB medium at an initial $OD_{600}$ of 0.005. Cells were grown at 37˚C until reaching exponential phase ($OD_{600}$ ~ 0.2 for *B. subtilis* and ~ 0.3 for *E. coli*), then subjected to cold shock by rapidly mixing the 37˚C culture with a defined volume of pre-chilled LB medium. For example, to achieve cold shock to 18˚C or 15˚C, 22 mL or 19 mL of the 37˚C culture was mixed with 20 mL of LB pre-chilled overnight at 4˚C, respectively. For cold shock to 12˚C, 13 mL of the 37˚C culture was mixed with 17 mL of ice-chilled medium; for 9˚C, 16mL of culture was mixed with 27 mL of the ice-chilled medium. After rapid mixing, cultures were immediately transferred to a water bath shaker pre-set to the target low temperature. Growth was monitored by measuring $OD_{600}$ every 20 minutes before cold shock, every 30 minutes during the first doubling time after cold shock, and every 1.5 hours thereafter. When $OD_{600}$ approached 0.3, cultures were back diluted to $OD_{600}$= 0.01 in pre-warmed or pre-chilled LB medium to prevent entry into stationary phase throughout the experiments.

#### Polysome Profiling

Strains were grown overnight at 37˚C and inoculated into 250 mL of LB at an initial $OD_{600}$ of 0.005. Pre- and post-cold shock samples were collected at designated time points. For 37˚C samples, 200 mL of culture was vacuum filtered through a 0.22 µm nitrocellulose membrane (GVS) in a 37˚C warm room. Post-cold shock samples at 18˚C were collected in the same manner, except at room temperature. Cell pellets were quickly scraped off the membrane using pre-warmed scoopula and flash-frozen in liquid nitrogen. Frozen pellets were stored at -80˚C until cell lysis.

Cell lysis was performed using 10 mL canisters (QIAGEN) pre-chilled in liquid nitrogen. Each cell pellet was lysed with 500 µL frozen lysis buffer (100mM $NH_4Cl$, 10mM $MgCl_2$, 5mM $CaCl_2$, 20mM Tris-HCl pH 8.0, 0.1% NP-40, 0.4% Triton X-100, 1mM chloramphenicol, 100 U/mL DNase I (Roche)). Lysis was carried out using a QIAGEN Tissue Lyser III for 5 cycles of 3 minutes at 15 Hz, with re-chilling the canisters in liquid nitrogen between cycles.

Pulverized lysates were thawed in a 30˚C water bath for 2 minutes and clarified by centrifugation at 16000 x g for 10 minutes at 4˚C. Clarified lysates containing 10 $A_{260}$ units were loaded onto 10–55% sucrose gradients prepared in gradient buffer (100mM NH4Cl, 10mM MgCl2, 5mM CaCl2, 20mM Tris-HCl pH 8.0, 100 µg/mL chloramphenicol, 2mM DTT) and ultracentrifuged at 35,000 rpm for 2.5 hours at 4˚C in an SW-41 Ti rotor (Beckman Coulter). Gradients were fractionated using a Biocomp Gradient Station (Biocomp Instruments), and polysome profiles were recorded by continuous $A_{260}$ monitoring with a TRIAX™ Flow Cell (Biocomp Instruments).

#### *Bacillus subtilis* Tn-seq

##### *B. subtilis Sample collection*

The *B. subtilis* transposon library was kindly provided by Alan Grossman (Massachusetts Institute of Technology) (9). Briefly, a library of random transposon insertions into *B. subtilis* genomic DNA was first made *in vitro* and then transformed into *B. subtilis* cells. A modified version of the *magellan6* transposon was used. Cells were plated on LB agar with spectinomycin, and colonies harboring transposon insertions were selected and pooled into a single library. This method generated approximately 3.85 x $10^5$ clones, each carrying a distinct transposon insertion (9).

For the cold shock phenotypic profiling, we first inoculated 200 µL of the frozen library into 20 mL LB media (initial $OD_{600}$ ~ 0.1) and grew the culture briefly to $OD_{600}$ ~ 0.4. A sample was collected as *Sample*

*#1* ($t_0$). The culture was then diluted to $OD_{600}$ ~ 0.02 in 50 mL LB media and grown at 37˚C with shaking at 250 rpm. A sample was collected at $OD_{600}$~ 0.3 as *Sample #2* (37˚C log phase). To induce cold shock, 25 mL of the 37˚C culture was mixed with 25 mL pre-chilled LB at 4˚C, resulting in an immediate temperature drop to 18˚C. The cultures were then quickly transferred to a water bath shaker maintained at 18˚C in a cold room. Samples were collected at 5 minutes (*Sample #3*, 18˚C 5 minutes) and 6 hours (*Sample #4*, 18˚C 1 doubling) after cold shock. To continue monitoring growth, the cultures were diluted to $OD_{600}$ ~ 0.01 in LB media pre-equilibrated to 18˚C and then grown overnight. Samples were collected the next day when $OD_{600}$ reached ~0.32 (*Sample #5*, 18˚C 6 doublings). Cultures were diluted again to $OD_{600}$ ~ 0.01 as described above, and another sample was collected the following day at $OD_{600}$ ~ 0.32 (*Sample #6*, 18˚C 11 doublings). For sample collection, cells (total $OD_{600}$ ~ 1.5) were harvested by centrifugation at 5000 x g for 10 minutes. Supernatants were discarded, and pellets were flash frozen in liquid nitrogen and stored at -80˚C.

***B. subtilis* *Tn-seq library construction and sequencing***

The procedure for constructing the Tn-seq library was modified based on previous works (9, 10). The inverted repeat sequences of the modified *magellan6* transposon contain an MmeI recognition site. MmeI cleaves DNA outside of its recognition sequence, resulting in 16 bp of genomic DNA attached to both ends of the transposon, which can be sequenced to map the transposon insertion sites.

To construct the sequencing libraries, genomic DNA (gDNA) was extracted from each sample using Qiagen DNeasy Blood and Tissue kit (Qiagen). 4 µg of purified gDNA was digested with the MmeI (NEB) at 37˚C for 2.5 hours. The 3' and 5' phosphates were removed by treatment with Quick CIP (NEB) at 37˚C for 20 min, followed by enzyme inactivation at 80˚C for 10 min. DNA fragments were cleaned up using the Zymo DNA Clean & Concentrator kit (Zymo Research) and eluted in 27.5 µL of 2mM Tris-Cl pH8.5.

A double-stranded DNA adapter was prepared by annealing oligos oYZ525 and oYZ526. Oligos were dissolved to 200 µM in 1mM Tris pH8.5. Equal volumes of the oligos were mixed, denatured at 96˚C for 2 min, and then allowed to cool to room temperature. The ligation reaction was prepared by mixing 27.5 µL of digested gDNA, 2 µL of annealed adapter, 3.5 µL of 10x T4 DNA ligase buffer, and 1.5 µL of T4 DNA ligase (NEB), followed by incubation overnight at 16˚C. DNA was then purified using the Qiagen PCR Purification kit and eluted with 35 µL EB buffer. 5 µL of the ligation product was used as template in a 50 µL PCR reaction with Q5 polymerase (NEB) for 16 cycles. Primers oYZ522 and oYZ523 were used, where NNNNNN denotes a 6-base index for multiplexing. Amplified libraries were concentrated using Zymo DNA Clean & Concentrator kit (Zymo Research) and ran on an 8% Invitrogen Novex TBE gel (Invitrogen). DNA bands of approximately 142 bp were excised and extracted to obtain the final sequencing library.

The constructed libraries were sequenced on an Illumina NovaSeq 6000 platform using the sequencing oligo oYZ524 to obtain the transposon-chromosome junction sequences, and the oligo oYZ527 to sequence the index barcode. After 16 sequencing cycles, raw data were collected from three different lanes and demultiplexed based on the index combinations. All the oligos used are listed in **Table S1**.

***B. subtilis* fitness value calculation**

Fitness values of individual genes were calculated as previously described (10), with modifications. After sequencing, 16-nt reads were mapped to the *B. subtilis* 168 genome (NC_000964.3) using Bowtie version 1.2.3 (5). Bowtie parameters were set to allow only reads that mapped to a unique genomic location without mismatches; all the other reads were excluded from downstream analysis. The number of Tn-seq reads mapped to each genomic location reflects the relative abundance of that transposon mutant in the pooled library.

Tn-seq read counts mapped to individual genomic locations were first summed across three technical replicates. Since transposon insertions do not distinguish between plus and minus strand, we combined reads mapped to either strand and generated wiggle files with two columns: genomic location and the corresponding Tn-seq read count. We then applied two normalization steps: (1) To normalize for chromosomal replication biases in Tn-seq read counts (11), the LOESS (Locally Estimated Scatterplot Smoothing) regression model was applied to fit a smooth curve through data points across the genome and adjusted the Tn-seq counts based on the prediction (**Fig. S2**). 2) To correct for the difference in total Tn-seq read counts across samples, Tn-seq data was normalized by the total number of mapped reads. The normalized Tn-seq data were then used to calculate the total read counts mapped to individual genes (i.e., $N_i(t)$ for gene *i* at time point *t*). Statistical analysis of normalized Tn-seq data, including cross-sample variance and significance testing, was performed using the R package edgeR (7). To ensure robustness, only genes with sufficient initial read counts (> 80; thresholds justified in **Fig. S5**) were analyzed.

We defined the fitness value of each gene as the relative growth advantage or disadvantage of its corresponding mutants compared to the rest of the pooled library over one cell generation. From time $t_1$ to $t_2$, the fitness of *gene i* mutants, denoted as $W_i$, was calulcated by the following equation:

$$W_i = \frac{\ln\,(N_i(t_2) \times \frac{d}{N_i(t_1)})}{ln((1 - N_i(t_2)) \times d/(1 - N_i(t_1)))}$$

Here, $N_i(t_1)$ and $N_i(t_2)$ represent the frequency of transposon mutants for gene *i* in the population at time point $t_1$ and $t_2$, respectively. $N_i(t)$ was calculated by summing the read counts mapped to the open reading frame of gene *i*, excluding the first 5% and last 10% of the gene, as transposon insertions in these regions are less likely to fully disrupt gene function. The expansion factor $d$ measures the fold increase in the total bacterial population between $t_1$ and $t_2$. Therefore, the fitness value is independent of the duration between time points, allowing comparison between stages of cold shock response and continuous growth. Normalized Tn-seq data and gene fitness values across all conditions are provided in **Tables S2–S3**.

#### *E. coli* RB (Random Barcode)-Tnseq

##### *E. coli Sample collection*

The *E. coli* BW25113 random-barcoded transposon mutant (RB-TnSeq) library was resourced from prior published work (12, 13). The frozen library was thawed completely and inoculated into 20 mL LB media (initial $OD_{600}$ ~ 0.1) and grown to $OD_{600}$ ~ 0.4. A sample was collected as *Sample #1* ($t_0$). The culture was then diluted to $OD_{600}$ ~ 0.02 in 50 mL LB media and grown at 37˚C to $OD_{600}$~ 0.3, collected as *Sample #2* (37˚C log phase). To induce cold shock, 25 mL of the 37˚C culture was mixed with 25 mL pre-chilled LB at 4˚C to reach 18˚C, or with 30 mL ice-cold LB to reach 15˚C. The cultures were quickly transferred to water bath shakers set at 18˚C or 15˚C. Samples were collected after 2 h at 18˚C or 4 h at 15˚C (*Sample #3*), and at 1, 6, and 11 doublings (*Samples #4–#*6) after cold shock. Cultures were diluted to $OD_{600}$ ~ 0.01 whenever $OD_{600}$ reached 0.3 to prevent entry into stationary phase.

##### *E. coli Bar-seq PCR and sequencing*

Genomic DNA was isolated from RB-TnSeq library samples using the DNeasy Blood and Tissue kit (Qiagen). We performed standard BarSeq PCR protocol as described previously (12, 13). Equal volumes (5 µL) of the individual BarSeq PCRs were pooled, and 50 µL of the pooled PCR product were purified with the DNA Clean and Concentrator kit (Zymo Research). The final BarSeq library was eluted in 40 µL of sterile water. The BarSeq libraries were sequenced on Illumina Novaseq instrument with 50 SE runs.

##### *E. coli fitness calculation*

Gene fitness was calculated as for *B. subtilis*. Chromosomal position bias was corrected using a LOESS model (**Figs. S3–S4**). Only genes with sufficient initial read counts (>40; thresholds justified in **Fig. S5**) were analyzed. Normalized Tn-seq data and gene fitness values across all conditions are provided in **Tables S4–S6**.

#### Supplementary Figures

**Figure S1**

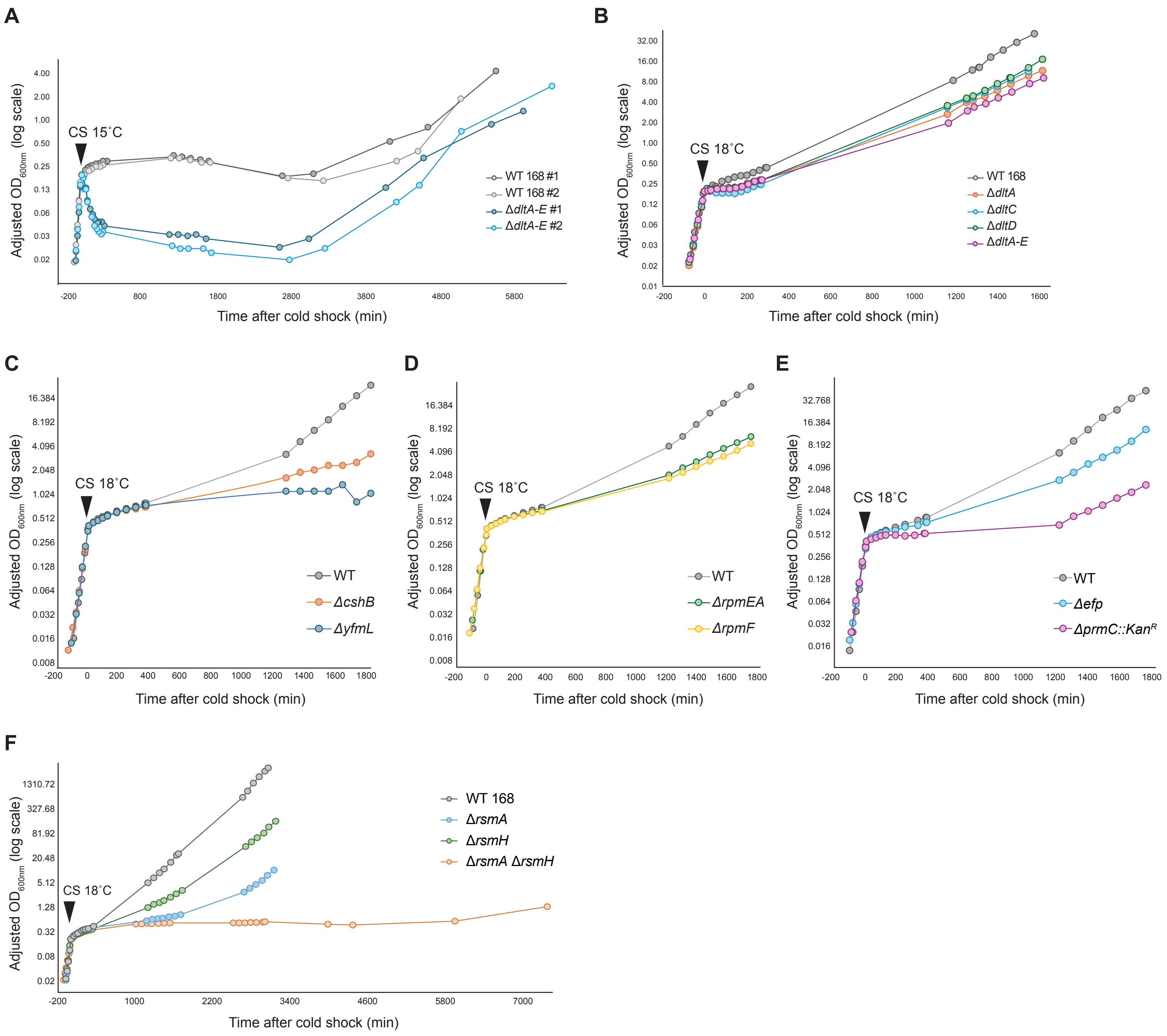
A
Adjusted OD600nm (log scale)
4.00
2.00
1.00
0.50
0.25
0.13
0.06
0.03
0.02
CS 15˚C
-200
800
1800
2800
3800
4800
5800
Time after cold shock (min)
WT 168 #1
WT 168 #2
ΔdltA-E #1
ΔdltA-E #2
B
Adjusted OD600nm (log scale)
32.00
16.00
8.00
4.00
2.00
1.00
0.50
0.25
0.12
0.06
0.03
0.02
0.01
CS 18˚C
-200
0
200
400
600
800
1000
1200
1400
1600
Time after cold shock (min)
WT 168
ΔdltA
ΔdltC
ΔdltD
ΔdltA-E
C
Adjusted OD600nm (log scale)
16.384
8.192
4.096
2.048
1.024
0.512
0.256
0.128
0.064
0.032
0.016
0.008
CS 18˚C
-200
0
200
400
600
800
1000
1200
1400
1600
1800
Time after cold shock (min)
WT
ΔcshB
ΔyfmL
D
Adjusted OD600nm (log scale)
16.384
8.192
4.096
2.048
1.024
0.512
0.256
0.128
0.064
0.032
0.016
0.008
CS 18˚C
-200
0
200
400
600
800
1000
1200
1400
1600
1800
Time after cold shock (min)
WT
ΔrpmEA
ΔrpmF
E
Adjusted OD600nm (log scale)
32.768
16.384
8.192
4.096
2.048
1.024
0.512
0.256
0.128
0.064
0.032
0.016
CS 18˚C
-200
0
200
400
600
800
1000
1200
1400
1600
1800
Time after cold shock (min)
WT
Δefp
ΔprmC::KanR
F
Adjusted OD600nm (log scale)
1310.72
327.68
81.92
20.48
5.12
1.28
0.32
0.08
0.02
CS 18˚C
-200
1000
2200
3400
4600
5800
7000
Time after cold shock (min)
WT 168
ΔrsmA
ΔrsmH
ΔrsmA ΔrsmH

**Figure S1. Growth phenotypes of *B. subtilis* mutants after cold shock to 18˚C**

**(A)** Growth curve of *B. subtilis* 168 (WT) and the Δ*dltA-E* mutant before and after cold shock. CS 15˚C: cold shock to 15˚C. Two biological replicates are shown as #1 and #2. X-axis: time after cold shock (min); Y-axis: adjusted optical density at 600nm ($OD_{600}$) on a log scale. $OD_{600}$ was adjusted at the transitions of cold shock and back-dilutions (see Methods) to allow continuous readings.

**(B)** Growth curve of WT, single *dlt* gene deletion mutants, and the Δ*dltA-E* mutant before and after cold shock to 18˚C.

**(C-E)** Growth curve of WT and various mutants before and after cold shock to 18˚C. The Δ*prmC::Kan*$^{R}$ mutant has a competence defect, making it challenging to construct a clean deletion; thus, a strain carrying the antibiotic marker was used.

**(F)** Growth curve of WT and various *rsmA* and *rsmH* mutants before and after cold shock to 18˚C. Δ*rsmA* Δ*rsmH* double mutant was cultured for additional days to monitor its long-term growth at 18˚C.

**Figure S2**

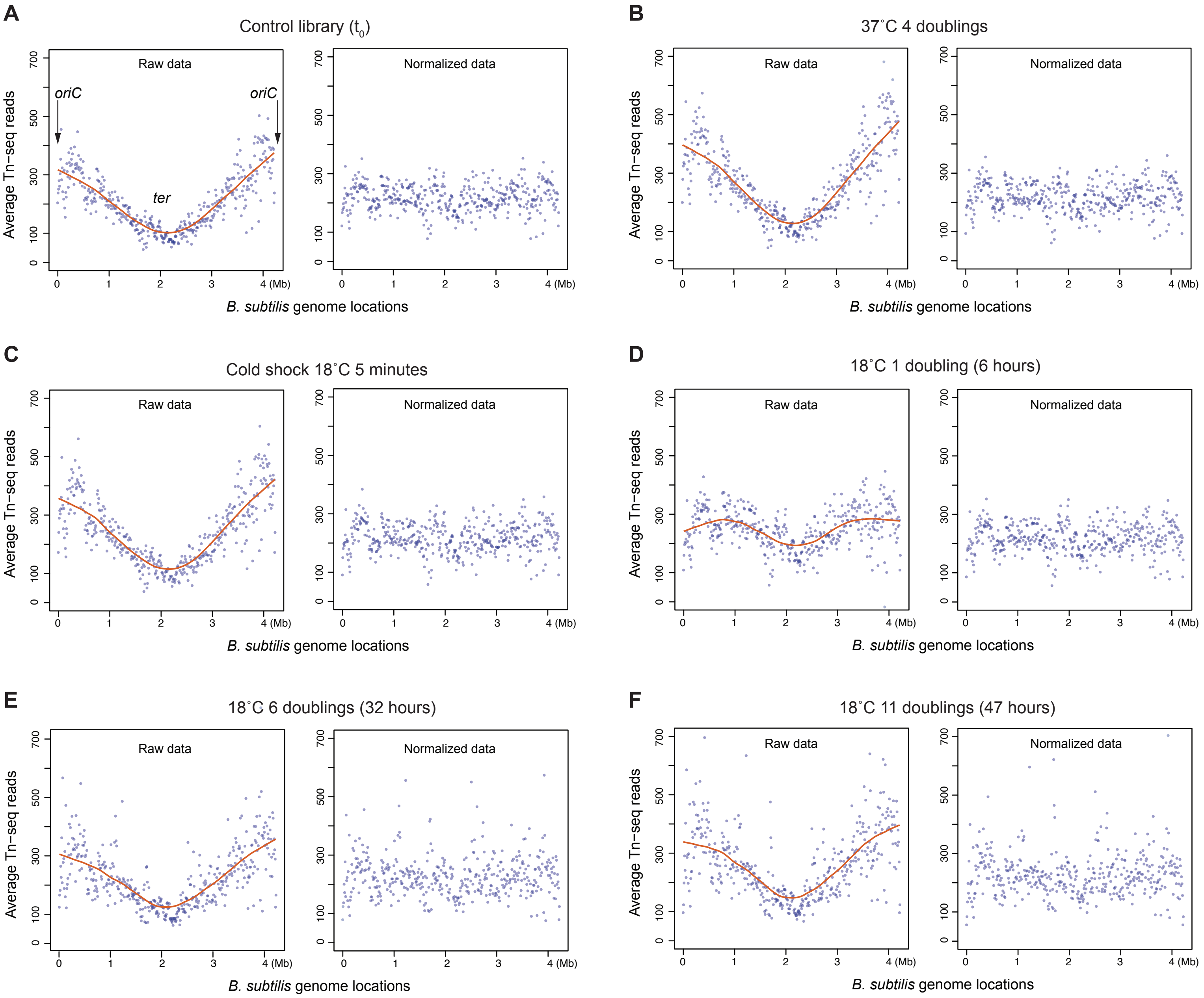
A
Control library ($t_0$)
Raw data
Normalized data
oriC
oriC
ter
Average Tn−seq reads
B. subtilis genome locations
0
100
300
500
700
0
1
2
3
4 (Mb)
B
37˚C 4 doublings
C
Cold shock 18˚C 5 minutes
D
18˚C 1 doubling (6 hours)
E
18˚C 6 doublings (32 hours)
F
18˚C 11 doublings (47 hours)

**Figure S2. Correction of chromosomal position bias in *B. subtilis* Tn-seq data using LOESS smoothing**

Comparison of raw and normalized Tn-seq data from *B. subtilis* samples collected at different time points: $t_0$ (control) **(A)**, 37˚C before cold shock **(B)**, and at 5 minutes **(C)**, or after 1, 6, or 11 doublings **(D-F)** following cold shock to 18˚C.

**Left panels:** Data points represent averaged Tn-seq read counts across the chromosome, calculated using a window size of 1/10,000 of the *B. subtilis* genome. Red curve: LOESS (Locally Estimated Scatterplot Smoothing) fit (span = 0.35, degree = 1). The origin of replication (*oriC*) is located at 0°/360° on the circular chromosome (14). *oriC* and the replication terminus region are labeled in **(A)**.

**Right panels:** Normalized Tn-seq read counts averaged across genomic windows. Normalization was performed by first correcting chromosomal position bias using the LOESS-predicted values, followed by scaling to the total number of mapped reads.

#### Figure S3

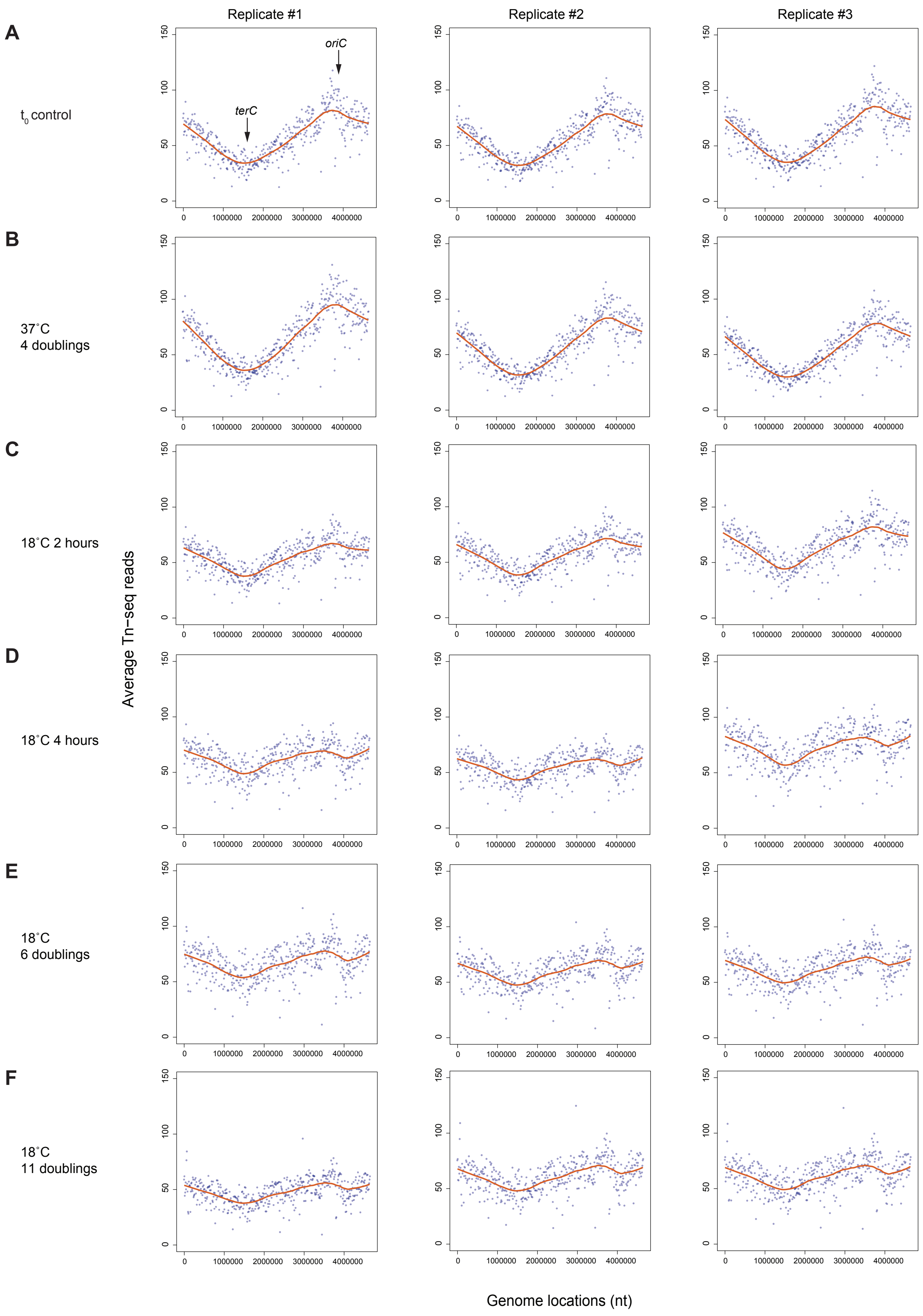

Replicate #1
Replicate #2
Replicate #3
A
t0 control
B
37˚C
4 doublings
C
18˚C 2 hours
D
18˚C 4 hours
E
18˚C
6 doublings
F
18˚C
11 doublings
oriC
terC
Average Tn−seq reads
Genome locations (nt)
0
50
100
150
0
1000000
2000000
3000000
4000000

**Figure S3. Correction of chromosomal position bias in *E. coli* Tn-seq data following cold shock at 18˚C using LOESS smoothing**

Averaged Tn-seq read counts across the *E. coli* chromosome, calculated using a window size of 1/10,000 of the genome. Red curve: LOESS fit (span = 0.25, degree = 1). The origin of replication (*oriC*), located at 304° on the circular chromosome, and the replication terminus site (*terC*) are labeled in **(A)**. Data are shown for three biological replicates collected at different time points: $t_0$ (control) **(A)**, 37˚C before cold shock **(B)**, and at 2 hours **(C)**, 4 hours **(D)**, or after 6, or 11 doublings **(E-F)** following cold shock to 18˚C.

#### Figure S4

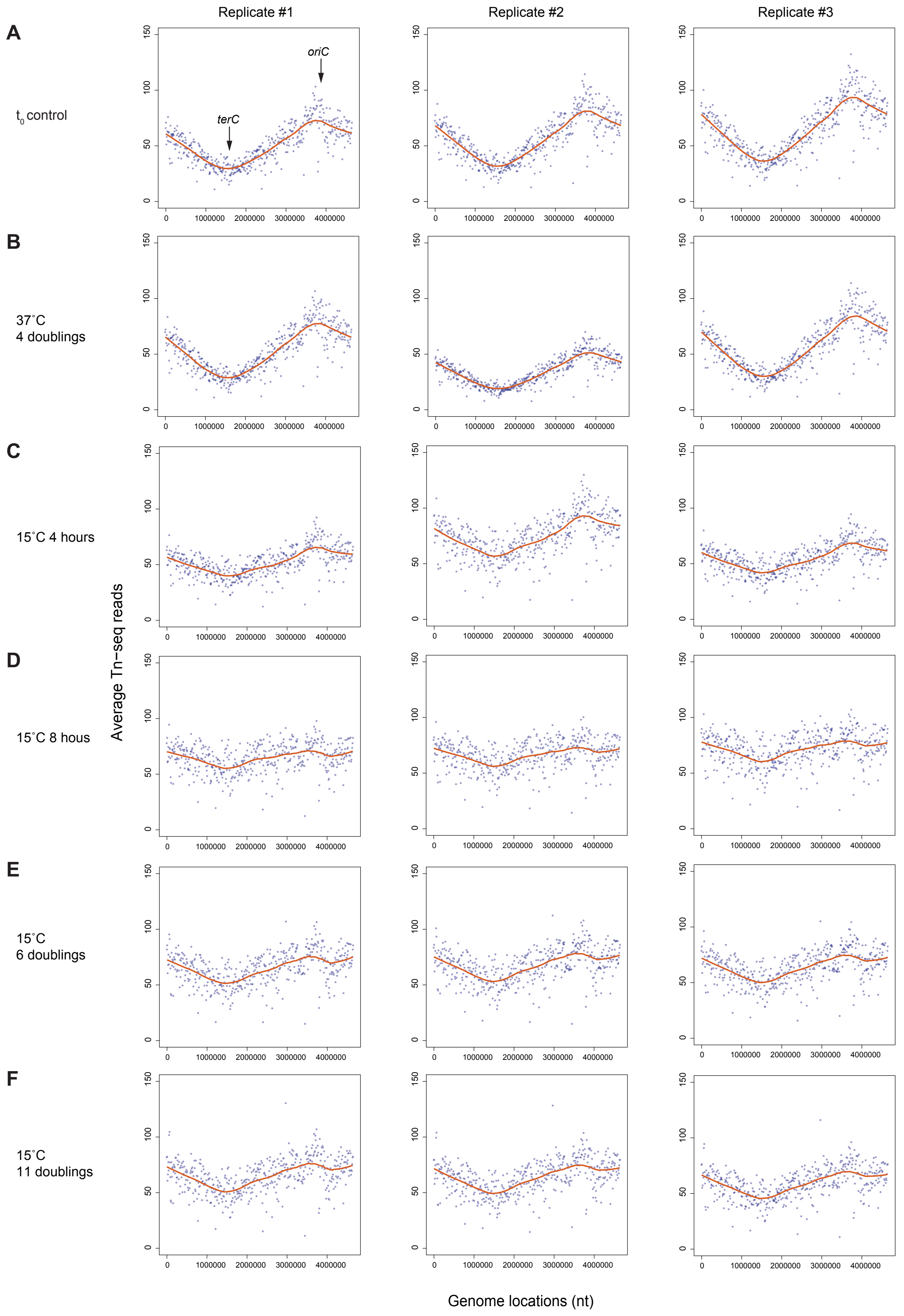
Replicate #1
Replicate #2
Replicate #3
A
t0 control
terC
oriC
B
37˚C
4 doublings
C
15˚C 4 hours
D
15˚C 8 hous
E
15˚C
6 doublings
F
15˚C
11 doublings
Average Tn−seq reads
Genome locations (nt)
0
50
100
150
1000000
2000000
3000000
4000000

**Figure S4. Correction of chromosomal position bias in *E. coli* Tn-seq data following cold shock at 15˚C using LOESS smoothing**

Averaged Tn-seq read counts across the *E. coli* chromosome, calculated using a window size of 1/10,000 of the genome. Red curve: LOESS fit (span = 0.25, degree = 1). The origin of replication (*oriC*), located at 304° on the circular chromosome, and the replication terminus site (*terC*) are labeled in **(A)**. Data are shown for three biological replicates collected at different time points: $t_0$ (control) **(A)**, 37˚C before cold shock **(B)**, and at 4 hours **(C)**, 8 hours **(D)**, or after 6, or 11 doublings **(E-F)** following cold shock to 15˚C.

### Figure S5

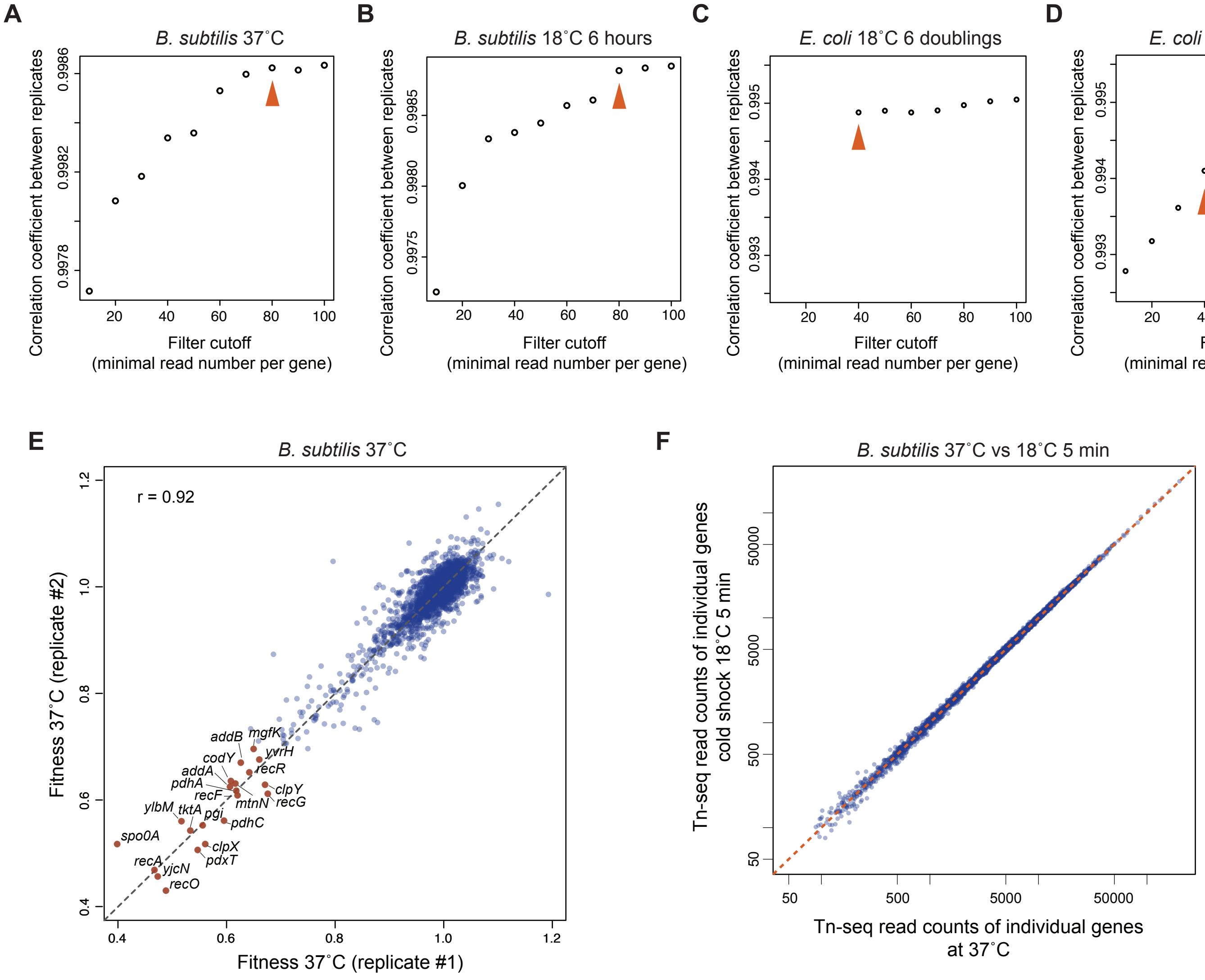

A
B. subtilis 37˚C
Correlation coefficient between replicates
Filter cutoff
(minimal read number per gene)
B
B. subtilis 18˚C 6 hours
C
E. coli 18˚C 6 doublings
D
E. coli 15˚C 6 doublings
E
B. subtilis 37˚C
r = 0.92
Fitness 37˚C (replicate #2)
Fitness 37˚C (replicate #1)
addB
mgfK
yvrH
codY
recR
addA
pdhA
clpY
recF
mtnN
recG
ylbM
tktA
pgi
pdhC
spo0A
clpX
pdxT
recA
yjcN
recO
F
B. subtilis 37˚C vs 18˚C 5 min
Tn-seq read counts of individual genes
cold shock 18˚C 5 min
Tn-seq read counts of individual genes
at 37˚C

**Figure S5. Gene filter determination and analysis of Tn-seq data at 37˚C**

**(A-D)** Determination of gene read count thresholds for inclusion in Tn-seq fitness analysis. Shown are example datasets from *B. subtilis* at 37˚C **(A)** and 6 hours after cold shock to 18˚C **(B)**, and *E. coli* 6 doublings after cold shock to 18˚C **(C)** or 15˚C **(D)**. X-axis: read count cutoff used as a filter to select genes with sufficient Tn-seq reads for fitness calculation; Y-axis: Person's correlation coefficient comparing gene-specific Tn-seq signals between two biological replicates, calculated using genes passing the filter. Final thresholds were chosen to maximize reproductivity while retaining a sufficient number of genes: 80 for *B. subtilis* and 40 for *E. coli* (shown by the red arrows).

**(E)** Scatter plot comparing gene-specific fitness values from two biological replicates of *B. subtilis* at 37˚C. 22 genes with fitness values < 0.7 are labeled. A corresponding set of *E. coli* genes is listed in Tables S5 and S6. r: Pearson's correlation coefficient; grey dashed lines: $y = x$.

**(F)** Scatter plot comparing gene-specific Tn-seq read counts in *B. subtilis* at 37˚C versus 5 minutes after cold shock to 18˚C. Red dashed lines: $y = x$.

### Figure S6

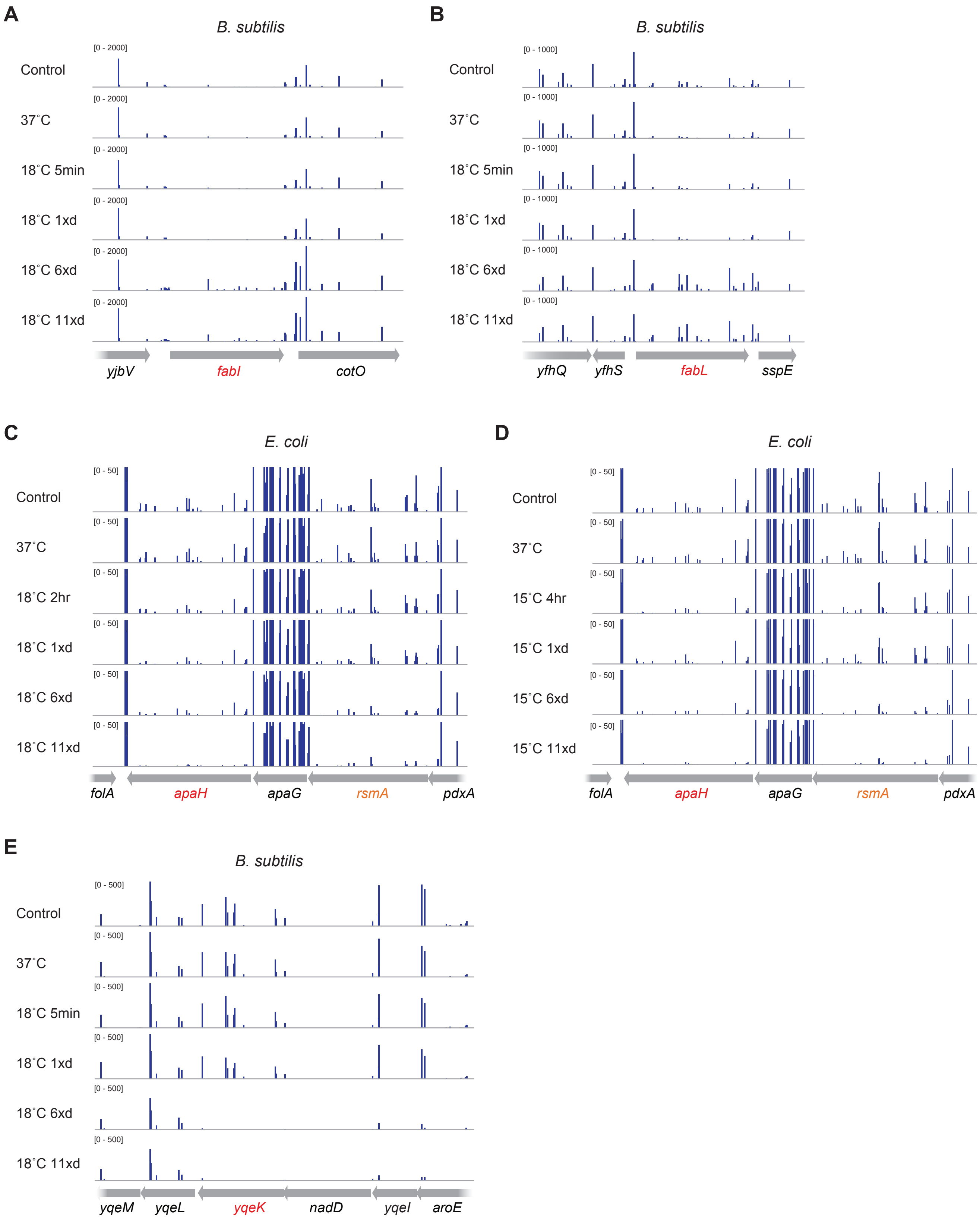
A
B. subtilis
[0 - 2000]
Control
37˚C
18˚C 5min
18˚C 1xd
18˚C 6xd
18˚C 11xd
yjbV
fabI
cotO
B
B. subtilis
[0 - 1000]
Control
37˚C
18˚C 5min
18˚C 1xd
18˚C 6xd
18˚C 11xd
yfhQ
yfhS
fabL
sspE
C
E. coli
[0 - 50]
Control
37˚C
18˚C 2hr
18˚C 1xd
18˚C 6xd
18˚C 11xd
folA
apaH
apaG
rsmA
pdxA
D
E. coli
[0 - 50]
Control
37˚C
15˚C 4hr
15˚C 1xd
15˚C 6xd
15˚C 11xd
folA
apaH
apaG
rsmA
pdxA
E
B. subtilis
[0 - 500]
Control
37˚C
18˚C 5min
18˚C 1xd
18˚C 6xd
18˚C 11xd
yqeM
yqeL
yqeK
nadD
yqeI
aroE

**Figure S6. Tn-seq data of key genes involved in the cold shock response**

**(A-B)** Normalized Tn-seq signals mapped to *fabI* **(A)**, *fabL* **(B)**, and their neighboring genes in *B. subtilis*, averaged from two biological replicates. Rows represent samples collected at $t_0$ (control), 37˚C before cold shock, and at 5 minutes, 1, 6, or 11 doublings after cold shock to 18˚C. Both *fabI* and *fabL* showed increased Tn-seq signals during the late stage of cold shock response.

**(C-E)** Normalized Tn-seq signals mapped to *apaH* in *E. coli* **(C, D)** and *yqeK* in *B. subtilis* **(E)**, along with their surrounding genomic regions. Signals were averaged from two or three biological replicates. For *E. coli*, samples were collected at $t_0$ (control), 37˚C, shortly after cold shock (2 hours at 18˚C or 4 hours at 15˚C), and at 1, 6, and 11 doublings after cold shock. For *B. subtilis*, the time points are the same as in panels **(A-B)**. Critical genes (*apaH* and *yqeK*) are highlighted in red, and *rsmA* is highlighted in orange.

#### Figure S7

**A**

| Doubling time (min) | 37˚C | 18˚C* |
|---|---|---|
| *B. subtilis 168* | 21.3 ± 1.0 | 190.8 ± 23.9 |
| Δ*rsmA* | 22.2 ± 1.5 | 359.5 ± 27.8 |
| Δ*rsmH* | 21.5 ± 1.6 | 258.5 ± 4.8 |
| Δ*rsmA* Δ*rsmH* | 30.9 ± 1.8 | 2961.6 ± 1221.0 |

**B**

| Doubling time (min) | 37˚C | 18˚C | 15˚C |
|---|---|---|---|
| WT MG1655 | 24.3 ± 1.1 | 198.6 ± 47.5 | 346.1 ± 58.2 |
| Δ*rsmA* | 26.5 ± 1.5 | 230.6 ± 41.1 | 450.0 ± 72.8 |
| Δ*rsmH* | 26.1 ± 2.5 | 226.9 ± 39.4 | 403.7 ± 81.8 |
| Δ*rsmA* Δ*rsmH* | 36.4 ± 1.1 | 344.1 ± 47.3 | 1341.5 ± 145.6 |

**C**

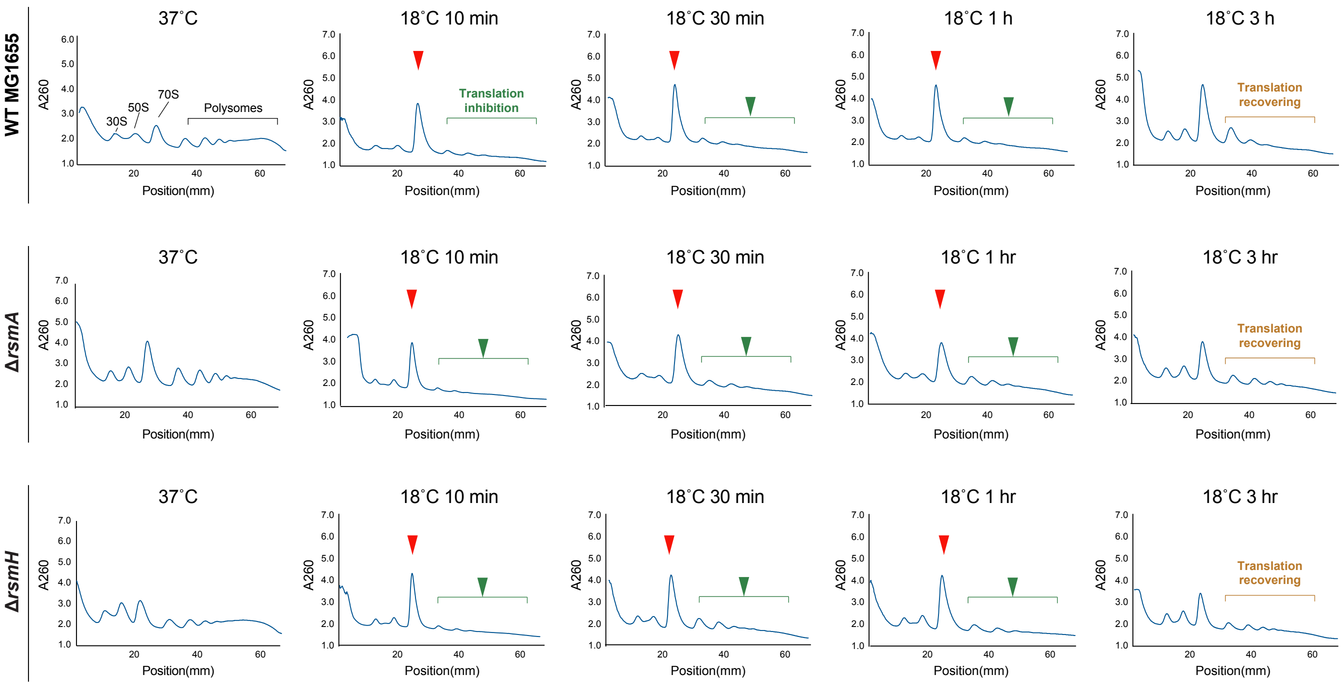

**Figure S7. Growth and polysome profiles of *rsmA, rsmH* mutants**

**(A-B)** Doubling times of *E. coli* **(A)** and *B. subtilis* **(B)** wild-type, Δ*rsmA*, Δ*rsmH,* and double mutants at 37˚C and after adaptation to 18˚C (*E. coli* and *B. subtilis*) or 15˚C (*E. coli*). For *B. subtilis*, the 18˚C doubling time was calculated from day 2. Data represent mean ± SD from at least three biological replicates.

**(C)** Polysome profiles of *E. coli* WT, Δ*rsmA,* and Δ*rsmH* mutants before and after cold shock. X-axis: position from the top of the centrifuge tube; Y-axis: UV absorbance at 260nm (A260). In the WT 37˚C profile, peaks corresponding to 30S and 50S ribosomal subunits, 70S monosomes, and polysomes are indicated. Cold shock induces accumulation of 70S monosomes (red arrows) and depletion of polysomes (green arrows). At 3 hours post-shock, translation shows partial recovery.

#### Supplementary Tables

**Table S1.** DNA oligos (Table S1.xlsx)
**Table S2.** *B. subtilis* Tn-seq read counts of individual genes (Table S2.xlsx)
**Table S3.** *B. subtilis* gene fitness 37˚C cold shock to 18˚C (Table S3.xlsx)
**Table S4.** *E. coli* Tn-seq read counts of individual genes (Table S4.xlsx)
**Table S5:** *E. coli* gene fitness 37˚C cold shock to 18˚C (Table S5.xlsx)
**Table S6:** *E. coli* gene fitness 37˚C cold shock to 15˚C (Table S6.xlsx)
